# Synthetic Nanobody-Functionalized Nanoparticles for Accelerated Development of Rapid, Accessible Detection of Viral Antigens

**DOI:** 10.1101/2021.05.09.443341

**Authors:** Xiahui Chen, Shoukai Kang, Ashif Ikbal, Zhi Zhao, Yuxin Pan, Jiawei Zuo, Liangcai Gu, Chao Wang

## Abstract

Successful control of emerging infectious diseases requires accelerated development of fast, affordable, and accessible assays to be widely implemented at a high frequency. Here we present a generalizable assay platform, <u>nano</u>body-functionalized <u>nano</u>particles for <u>r</u>apid, <u>e</u>lectronic <u>d</u>etection (Nano2RED), demonstrated in the detection of Ebola and COVID-19 antigens. To efficiently generate high-quality affinity reagents, synthetic nanobody co-binders and mono-binders with high affinity, specificity, and stability were selected by phage display screening of a vastly diverse, rationally randomized combinatorial library, bacterially expressed and site-specifically conjugated to gold nanoparticles (AuNPs) as multivalent in-solution sensors. Without requiring fluorescent labelling, washing, or enzymatic amplification, these AuNPs reliably transduce antigen binding signals upon mixing into physical AuNP aggregation and sedimentation processes, displaying antigen-dependent optical extinction readily detectable by spectrometry or simple electronic circuitry. With nanobodies against an Ebola virus secreted glycoprotein (sGP) and a SARS-CoV-2 spike protein receptor binding domain (RBD) as targets, Nano2RED showed a high sensitivity (limit of detection of ∼10 pg/mL for sGP and ∼40 pg/mL for RBD in diluted human serum), a high specificity, and a large dynamic range (∼7 logs). Unlike conventional assays where slow mass transport for surface binding limits the assay time, Nano2RED features fast antigen diffusion at micrometer scale, and can be accelerated to deliver results within a few minutes. The rapid detection, low material cost (estimated < $0.01 per test), inexpensive and portable readout system (< $5 and < 100 cm^3^), and digital data output, make Nano2RED particularly suitable for screening of patient samples with simplified operation and accelerated data transmission. Our method is widely applicable for prototyping diagnostic assays for other antigens from new emerging viruses.

## Introduction

In recent times, we have witnessed the emergence of many infectious viral diseases, from the highly fatal Ebola virus disease (EVD, with a fatality rate of 45% to 90%)^1, 2^ to the highly contagious coronavirus disease 2019 (COVID-19) with its global >200 million infections and >4 million deaths as of August 2021^3^. Future emergence of Disease X, as contagious as COVID-19 and as lethal as EVD, would pose an even greater threat to humanity, and will be both difficult to prevent or predict. During disease emergence, early pathogen identification and infection isolation are extremely critical for containing disease transmission^4^. Therefore, for effective mitigation, it is necessary to accelerate the design, development, and validation of diagnostic processes, as well as to make the diagnostic tools broadly accessible within weeks of the initial outbreak^4^.

Current diagnostic methods rely on the detection of the genetic (or molecular), antigenic, or serological (antibody) markers^5^. Genetic diagnostics use DNA sequencing, polymerase amplification assays, or most recently, CRISPR technologies^6, 7^. For example, real-time reverse transcription polymerase chain reaction (RT-PCR) tests are viewed as the gold standard for their high sensitivity; however, these tests are also costly, time-consuming, and instrument-heavy. Genetic tests also can often display false positives by picking up genetic fragments from inactive viruses^8^. In comparison, antigen and antibody detections are complementary as they allow more rapid, affordable, and accessible detection without complex sample preparation or amplification. As such, these detection methods are viewed as suitable for surveillance and timely isolation of highly infectious individuals, particularly outside clinical settings^8^. While antibody (*e*.*g*., IgM) detection has been used for disease diagnostics^9^, it is less predictive and more suitable for immune response studies. In comparison, viral protein antigen tests provide a reliable field-test solution in diagnosing symptomatic patients^9-11^, and may serve to screen asymptomatic contacts that may become symptomatic^12^. In addition, since they are rapid, easy to operate, and low-cost, antigen tests can be deployed at high frequencies and large volumes for in-time surveillance, which is thought to be the most important factor in disrupting a virus transmission chain^8, 12^.

Current antigen diagnostics typically employ enzyme-linked immunosorbent assays (ELISA) and lateral flow immunoassays (LFIs). ELISA is the workhorse for analyzing antigens and antibodies, but it requires a multistep workflow and a series of washing steps, hours of incubation prior to readout, and a readout system dependent on substrate conversion and luminescence recording. Deployment of ELISAs in high-throughput mass screenings requires automated liquid handling systems to coordinate the complex workflow, which is not ideal for portable uses. LFIs are potentially much easier to use outside lab settings but usually have much lower sensitivity and thus poorer accuracy compared to ELISA^13^.

Here we report a modular strategy, i.e., <u>nano</u>body-conjugated <u>nano</u>particles for <u>r</u>apid <u>e</u>lectronic <u>d</u>etection (Nano2RED), which can quickly establish a rapid, accessible antigen diagnostic tool within a few weeks of pathogen identification. To generate high-quality affinity reagents within two weeks for any given purified marker protein, we have streamlined a protocol for phage selection of single-domain antibodies (or nanobodies)^14^ from a highly diverse combinatorial library (> 10^9^) and bacterial expression of top hits with a significantly faster turnaround time and a lower cost than mammalian expression. Unlike traditional antibodies requiring non-specific conjugation to any solid support, potentially resulting in loss of function, nanobodies were genetically fused with an AviTag for site-specific biotinylation and immobilization onto streptavidin-coated gold nanoparticles (AuNPs). These nanobody-functionalized AuNPs serve as multivalent antigen binding sensors in our Nano2RED assays for Ebola and COVID-19 antigen detection. Based on only physical processes, including plasmonic NP color display, NP precipitation, and semiconductor photon absorption, Nano2RED quantitatively transduces antigen binding into colorimetric, spectrometric, and electronic readouts (Figure. 1). Without fluorescent labeling or chemiluminescent readout, Nano2RED differs fundamentally from conventional high-sensitivity tests (e.g., genetic tests or ELISA) that are generally expensive and more suitable for lab use. Yet, Nano2RED greatly outperforms conventional portable and low-cost tests (e.g., LFIs), which are not qualitative or sensitive enough. Uniquely, Nano2RED features portability, low cost, and simplicity while preserving a high sensitivity (LOD of ∼ 0.13 pM or 11 pg/mL in Ebola sGP sensing), a high specificity (distinguishing sGP from its membrane-anchored isoform GP1,2) and a large dynamic range (∼7 logs). Additionally, its electronic readout capability can be extended to automate data collection, storage, and analysis, further reducing the workload health care workers, and speeding up diagnostic and surveillance response.

**Figure 1.**
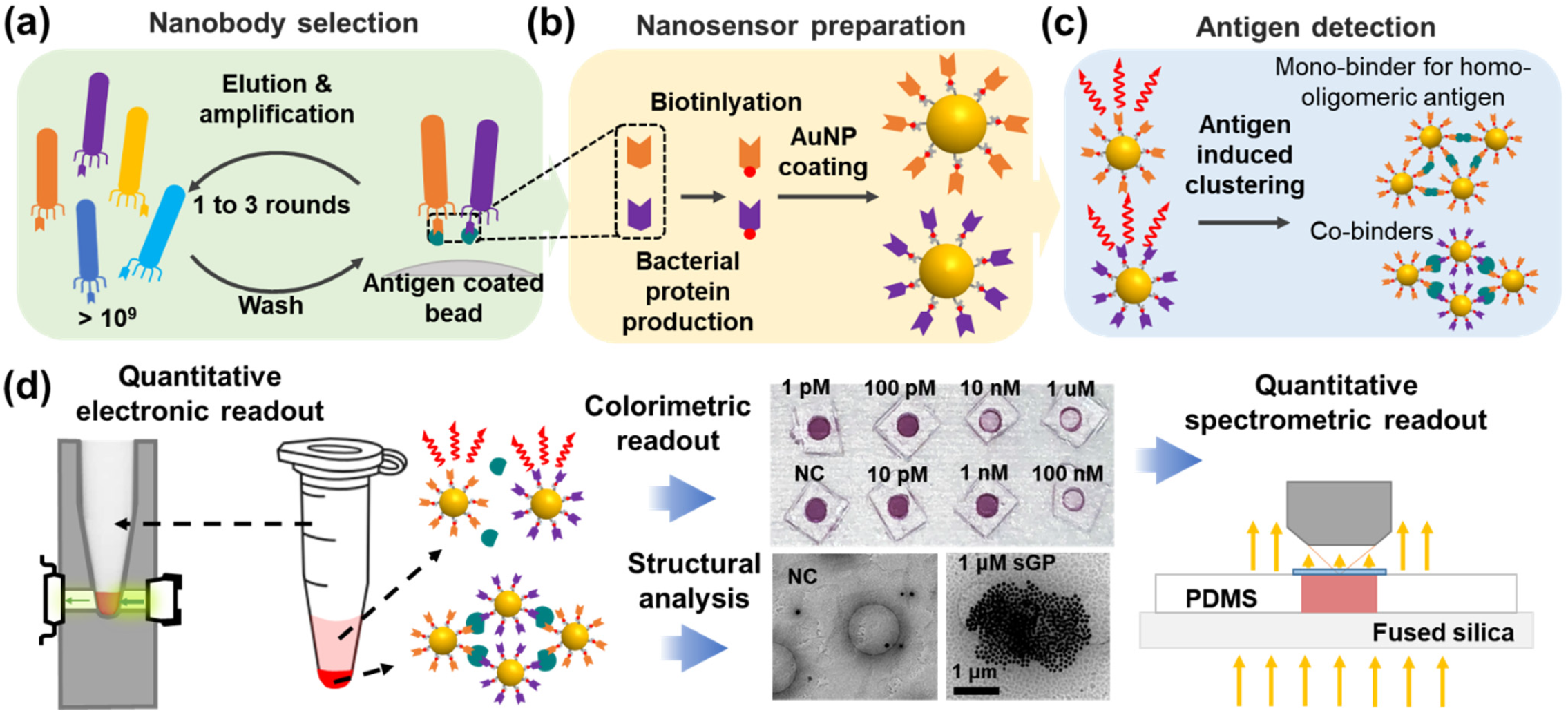
Overview of Nano2RED assay development and characterizations. (a)-(c) Key steps of nanobody selection and surface function of nanobody on AuNPs. (d) Schematics showing different characterization and readout methods for understanding the assay mechanism, and colorimetric and quantitative determination of antigen concentration.

### Nanobody co-binder selection for AuNP functionalization

We generated nanobody co-binders (*i*.*e*., two mono-binders simultaneously binding to non-overlapping epitopes in the same antigen) against target antigens for a new in-solution assay to improve the sensitivity and specificity (Figure 1**a**). Traditional methods for selecting antibody-based co-binders are slow and costly, so here we established a fast, robust protocol including the phage display selection of the combinatorial nanobody library, parallel bacterial protein production, co-binder validation, and AuNP functionalization that can be completed in less than two weeks upon the availability of an antigen protein (Figure 2**a**). Nanobodies, a single-domain (12-15 kDa) functional antibody fragments from camelid comprising a universal scaffold and three variable complementarity-determining regions, are ideally suited for phage display selection and low-cost bacterial production^14^. To avoid relatively lengthy and costly procedures and animal protection issues associated with traditional antibody screening, we screened the synthetic nanobodies library with an optimized thermostable scaffold prepared in our previous work^15^. The Ebola antigen, sGP, is a homodimeric isoform of the glycoproteins encoded by a GP gene of all five species of *Ebolavirus* with multiple post-translational modificiations^16^. sGP is believed to act as a decoy to disrupt the host immune system by absorbing anti-GP antibodies^16, 17^. Given its abundance in the blood stream upon infection and its quantitative correlation with disease progression and humoral response, sGP is widely used as a circulating biomarker in EBOV diagnostics^17, 18^. The chosen SARS-CoV-2 antigen, RBD, engages the viral receptor, human angiotensin-converting enzyme 2 (ACE2), causing conformational changes that trigger a cleavage needed for viral infection^19, 20^. It is a major antigenic target for protective antibodies^21^, and thus is highly significant for diagnostics, as well as for the development of vaccines and therapeutic neutralizing antibodies^22, 23^.

**Figure 2.**
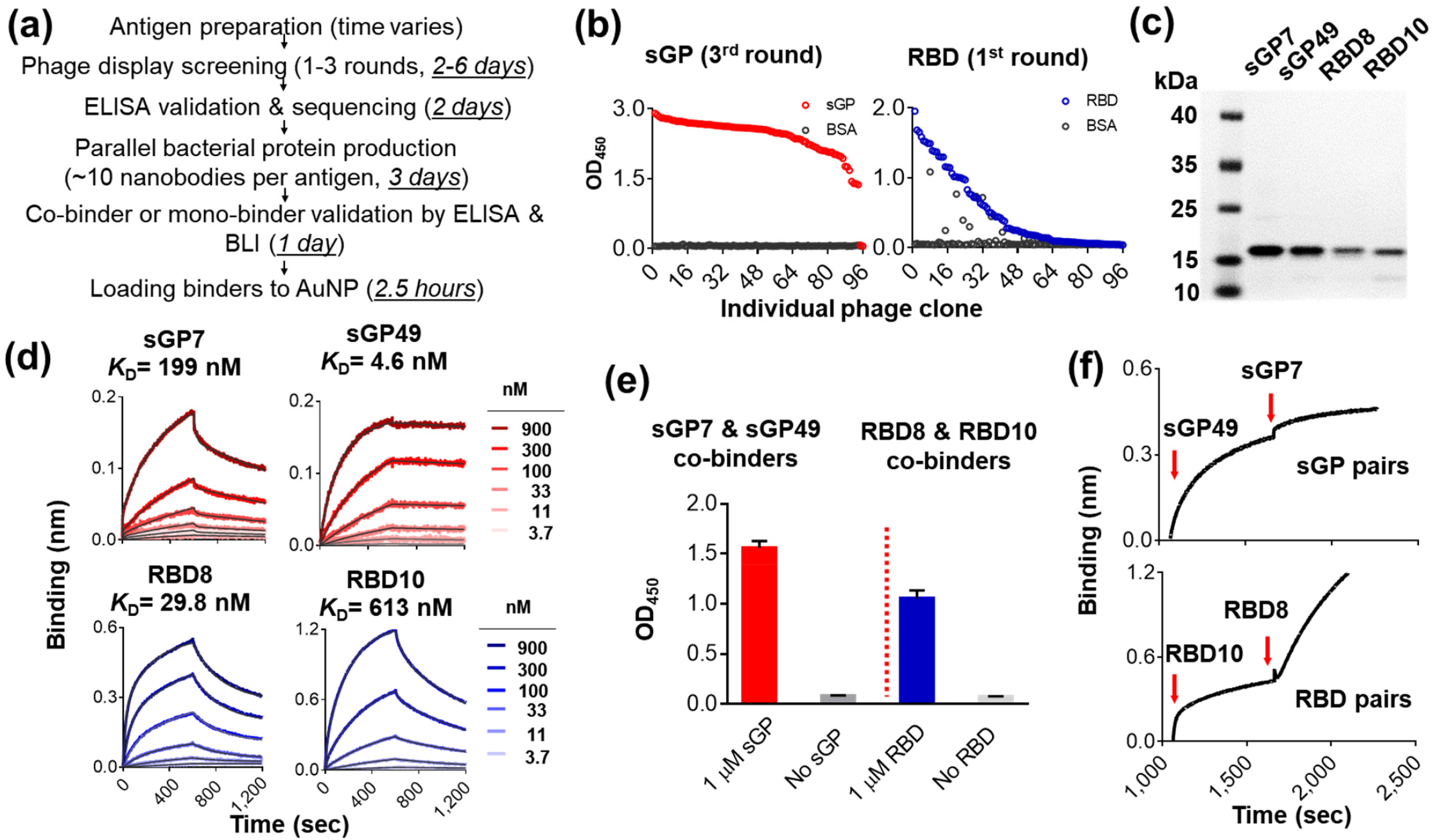
Identification and characterization of antigen-specific nanobody binders. (a) The flowchart and timeline of nanobody binder identification. (b) Binding of single phage clones to antigen measured by ELISA. Biotinylated sGP (red) and RBD (blue) were immobilized on streptavidin-coated plates, respectively. BSA was used as a control. (c) SDS-PAGE analysis of purified sGP and RBD-specific binders. (d) BLI analysis of specific binders binding to sGP (upper: red lines) and RBD (lower: blue lines) at different concentrations. The sGP and RBD were immobilized on streptavidin biosensors (SA). Measured data were globally fitted (grey lines). (e) Co-binder validation by sandwich ELISA. The first biotinylated nanobody binders (sGP 49 and RBD 10) were immobilized on a plate, incubated with or without antigen, and then the second binders (sGP7 and RBD8) were detected by a horseradish peroxidase-conjugated antibody. (f) Co-binder validation by two-step binding characterization using BLI. Biotinylated sGP (top panel) and RBD (bottom panel) were immobilized on SA sensors. Epitope binning was performed by first dipping into the first binder well for 750 s for saturation and then incubation with second binder.

To efficiently identify nanobodies that can bind non-overlapping epitopes of an antigen protein, we assessed clonal diversity and co-binding abilities of candidates enriched in different biopanning rounds. The antigens, sGP (ManoRiver) and RBD (residues 328-531), were expressed as AviTag fusions in HEK293 and biotinylated for immobilization on streptavidin-coated magnetic beads. We identified three co-binder pairs from 10 unique nanobodies out of 96 randomly picked clones that specifically bind to sGP after three rounds of biopanning. For the RBD with a relatively smaller size (∼30 kDa), we identified two co-binder pairs from 12 unique binders after the first round (Figure 2b). The two top co-binder pairs, termed sGP7–sGP49 and RBD8–RBD10, were bacterially expressed and purified (Figure 2c and Figure S1a) with high yields (1.5 to 6 mg per liter of culture). Their equilibrium dissociation constants (*K*_*D*_) were measured to be in the nanomolar range by Bio-Layer Interferometry (BLI) (Figure 2**d** and Figure S1 **a**) and the co-binding activities were validated by ELISA (Figure 2**e** and Figure S1 **b**) and BLI (Figure 2**f** and Figure S1 **b**). Lastly, nanobodies were biotinylated with *E. coli* biotin ligase (BirA) as previously reported^15^ and then loaded to streptavidin-coated AuNPs (see Methods section).

### Nanobody-functionalized nanoparticles for sensing

In our assay design, AuNPs densely coated with biotinylated nanobodies allow multivalent antigen sensing (Figure 1**a** and 1**b**) known to significantly enhance antigen binding compared to the monovalent binding^24, 25^. Further, the multivalent binding also facilitates AuNP aggregation at the presence of the antigen and subsequent precipitation, producing antigen-concentration-dependent signals within minutes. The AuNP aggregation is further quantified by optical and electronic measures. In our proposed sensing scheme (Figure 1**b**), AuNPs, without nonspecific particle-particle interaction, are initially homogenously dispersed in colloid, presenting a reddish color from characteristic localized surface plasmon resonance (LSPR) extinction^26^. Upon mixing with viral antigens, multiple AuNPs are pulled together by the antigen-nanobody binding to gradually form large aggregates. Compared to a single AuNP, the formation of AuNP aggregates gradually shifts LSPR extinction to higher wavelengths with broadened resonance attributed to plasmonic coupling between AuNPs^27, 28^, a phenomenon that can be simulated by finite-difference time-domain (FDTD) method (Supplementary section 2 and Figure S2). This leads to increased transparency of the AuNP colloid preciously described in DNA and protein sensing applications^29, 30^. Large AuNP aggregates can form pellets as gravity overtakes the fluidic drag force (Figure 1**c-d**). As a result, decreased AuNP concentrations in the upper liquid result in a colorimetric change correlated with sGP concentrations (Figure 1**d**). The color change can be directly visualized by eye, and quantified in a well plate by spectrophotometer or using a simple electronic device that measures the AuNP extinction.

### Colorimetric and spectrometric sensing of sGP

The size and shape of AuNPs determine the optical extinction and therefore the suspension color, hence affecting the sensitivity and assay incubation time (Figures S3 and S4). Here, the AuNP size effect was studied with NP diameters of 40, 60, 80, and 100 nm in sensing of Ebola sGP proteins using sGP49 nanobody in 1× phosphate buffered saline (PBS) buffer (Supplementary section 3). To standardize the measurement, the sGP signals were collected using a UV-visible spectrometer coupled to an upright microscope (Figure S3**a**). We custom-designed a polydimethylsiloxane (PDMS) well plate bonded to glass slides as the sample cuvette (Figure S3**b**). Top-level liquid from sGP sensing samples (5 μL) after incubation were loaded and inspected by optical imaging and spectroscopy readout. Clearly, as evidenced in the optical images (Figure S4 **a-d**), the color of the assay is redder for small NPs but greener for larger ones, which is attributed to a redshift in extinction resonance wavelengths at larger NP sizes. Additionally, a significant color contrast was observed in distinguishing 10 nM and higher sGP concentration from the reference negative control (NC) sample (with only PBS buffer but no sGP) for all AuNP sizes, indicating that sGP can be readily detected by the naked eye. Such colorimetric diagnostics would be very useful for qualitative or semi-quantitative diagnostics in resource-limited settings, but less ideal for quantitative and ultrasensitive detection.

Additional accurate sGP detection was performed by quantifying the AuNP extinction signals in the PDMS plate using our spectroscopic system (Figure S4 **e-l**). The AuNP extinction is correlated with its concentration [*NP*] and diameter *d* following *σ*_*ext*_ *∝* [*NP*]*d*^3 31^. A decrease of extinction indicated a drop of [*NP*] in the upper liquid level caused by antigen-induced AuNP precipitation. Further, the AuNP extinction peak values were extracted and plotted as standard curves against the sGP concentration at each AuNP size (Figure S4 **i-l**). This incubation-based assay had a large dynamic range of ∼100 pM to ∼100 nM for all AuNP sizes. In addition, the incubation was found to take 4 to 7 hours, using 40 to 100 nm NPs for detecting 10 nM sGP in 1× PBS (Figure S4 **m-p**). This NP size-dependent response could be understood intuitively from the antigen binding dynamics and the AuNP precipitation process. On the one hand, smaller NPs had higher starting concentrations, given that in our design the starting suspension extinction *σ*_*ext*_ *∝* [*NP*]*d*^3^ was about the same for all sizes, and therefore were expected to initiate the antigen-binding and NP aggregation reaction relatively faster. On the other hand, the precipitation of smaller aggregates took a longer time, resulting in a longer incubation period. From these experimental analyses in 1× PBS buffer, we chose 80 nm AuNPs to further characterize the assay performance in sGP sensing (Figure 3**a**). This selection was based on several factors: their slightly higher sensitivity (∼15 pM, compared to ∼100 pM for other sizes), larger detection dynamic range (up to 4 logs, compared to 2 to 3 logs for other sizes), and shorter incubation time (3 to 4 hours, compared to 4 to 7 hours for other sizes).

**Figure 3.**
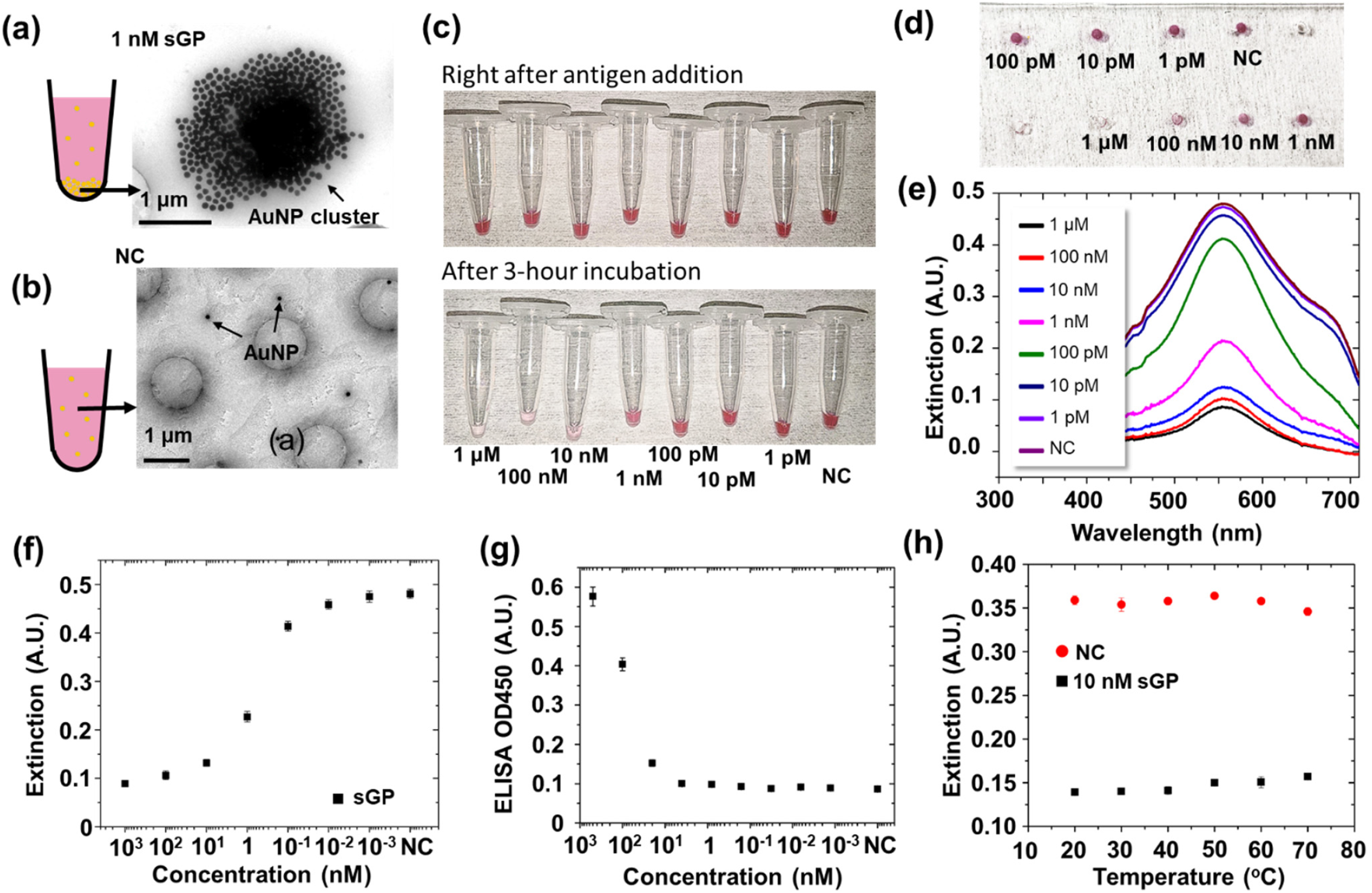
Ebola sGP sensing using mono-binder antibody sGP49 by incubation. (a-b) Cryo-TEM image of precipitates after 3-h incubation: (a) with 1 nM sGP, (b) the negative control (NC) sample (no sGP). (c) Visual images of samples loaded in microcentrifuge tubes, right after mixing and 3 hours after incubation. (d) The upper-level liquid samples loaded in PDMS well plate. (e) Extinction spectra measured from PDMS well plate. (f) Extinction peak (559 nm) values plotted against sGP and GP1,2 concentrations. (g) Optical signals (optical density at 450 nm) measured by sandwich ELISA in detection of sGP. (h) Extinction peak (559 nm) values extracted from spectroscopic measurements in detecting 10 nM sGP and NC at temperatures from 20 °C to 70 °C. The buffer was 1 × PBS in Figures a and b, and 5% FBS in Figures c to h. The sGP concentration was from 1 pM to 1 μM. NC sample was the buffer without sGP or GP1,2. The AuNPs were 80 nm in all measurements.

To understand the assay’s working mechanism, we complemented the solution-phase optical testing by inspecting the AuNP precipitates in solid state using different structural and optical characterization methods (Supplementary section 4). First, cryogenic transmission electron microscope (CryoTEM) images showed aggregates of AuNPs formed with 1 nM sGP (Figure 3**a**) with an average cluster size of 1.8 by 1.4 μm, while only 80 nm AuNP without clusters were observed in the upper-level liquid (Figure 3**b**) or in the precipitates of the NC sample (Figure S5). This supports our sensing mechanism in that AuNP precipitation serves to transduce antigen binding to solution color change for sensing readout (Figure 1). Further, we have performed drop-casting to deposit AuNP upper-level liquid samples on glass slides for optical extinction analysis (Figures S6 and S7) and on gold films for scanning electron microscopy (SEM) and dark field scattering imaging (Figures S8, S9, and S10). The measurement results were, in general, consistent with spectrometric in-solution sGP detection using PDMS well plate, but inferior in sensitivity (150 pM for SEM, ∼174 pM for drop-cast on glass slide, and ∼1 nM for dark filed imaging, Table 1). The decreased sensitivity could be attributed to inherent variations associated with sample preparation and background noise in the readout systems. These solid-state characterization methods were non-ideal for accessible and precise detection, given the low sensitivity and need of lab instrument for readout, but they provided valuable insight into the nano-scale NP aggregation process.

**Table 1.**
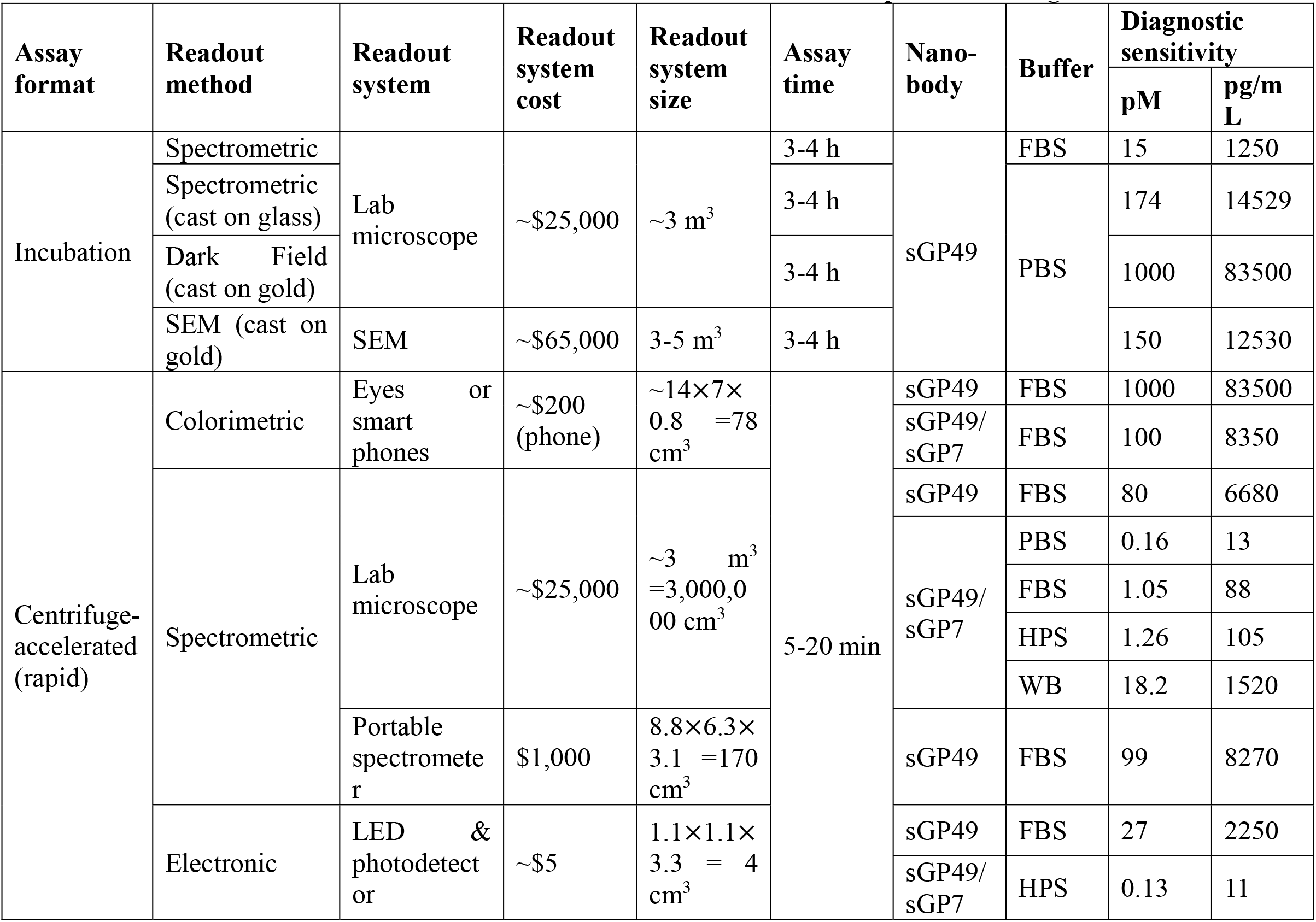
Performance of the Nano2RED in Ebola sGP protein sensing.

sGP was further detected in diluted fetal bovine serum (FBS, 5%) using 80 nm sGP49-functionalized AuNPs. Similarly, to test in 1× PBS, after 3-hour incubation in microcentrifuge tubes (Figure 3**c**), the upper-level liquid samples were loaded into a PDMS well plate (Figure 3**d**) and measured by spectrometer (Figure 3**e**). From the plot of extinction peak values against sGP concentration (Figure 3**f**), our assay could again detect sGP over a broad range from 10 pM to 100 nM, which supports clinically relevant Ebola detection from patients’ blood (sub-nM to μM) ^32, 33^. Here the three-sigma limit of detection (LoD), defined as the concentration displaying an extinction differentiable from the NC sample (*E*_*NC*_), or *E*_*NC*_ - 3*σ* where *σ* is the measurement variation of all samples, was found to be about 15 pM (or 1.25 ng/mL), comparable to that measured using sGP49 phage ELISA (LOD estimated ∼80 pM, Figure 3**g** and supplementary Table S1). The LOD can be understood from simple and rough estimations based on the nature of multivalent antigen binding (supplementary section 5). We also found the 10 nM sGP could be easily distinguished from NC sample at a broad temperature range from 20 to 70°C (Figure 3**h**). This indicates our assay can be transported, stored, and tested at ambient temperatures without serious concerns of performance degradation, which is very important for mass screening.

### Rapid antigen detection

The PDMS well plate-based spectrometric measurement required about 3 hours incubation for effective AuNP bridging and precipitation, which is shorter than ELISA and much better than many RT-PCR assays. However, rapid diagnostics, that is, less than 30 minutes, is more desirable for accessible infectious disease diagnosis and control of disease spread. Here, we further studied the sensing mechanism, aiming to reduce the detection time (Supplementary section 6.1). In conventional ELISA assays, the antigen diffusion process is usually the rate-limiting step^34^, since the fluidic transport to a solid surface is ineffective given long diffusion length (millimeter scale liquid depth in well plate) and slow fluidic flow speed at plane surfaces (near zero surface velocity). This leads to slow mass transport and ineffective surface binding, and an accordingly long assay time. Differently, diffusion is no longer the limiting factor in our assay, given the use of NPs in lieu of plane surfaces as reaction sites. The diffusion length is estimated to be about 1 μm due to a high NP concentration (*e*.*g*. ∼0.036 nM for 80 nm AuNPs). The small size and mass of AuNPs (5 × 10^−15^*g*) result in a high diffusivity (*D*_*NP*_ ∼4.2 μm^2^/s from Stokes*−*Einstein equation) and a high thermal velocity (estimated 0.028 m/s). All of these features promote effective fluidic transport and antigen binding, with an estimated diffusion time of <1 sec.

We further speculated that the AuNP aggregation and precipitation process could play important roles in determining assay time. Here, we developed a simplified model based on Smoluchowski’s coagulation equation to understand the aggregation process (Figure 4 **a-b**, and more details in Supplementary section 6.2)^35^. Briefly, an empirical parameter *P*, which defines the probability of antigen-nanobody binding per collision, has a large impact on the modelled assay time. By comparing to experimentally measured assay incubation time, we found using *P*=1, that is very high-affinity binding, provides a much better prediction compared to using a smaller *P* value calculated by the ELISA-measured kinetic constants. This observation is attributed to two factors: the ELISA-measured binding kinetics is strongly affected by the surface-limited diffusion process and could not precisely estimate the true nanobody-antigen binding in solution^36, 37^; and the multivalence of the nanobody-bound AuNPs greatly improves the observed “functional affinity” compared to intrinsic mono-binding affinity ^24, 25^. Using this model, it was estimated that the aggregation time constant *τ*_*agg*_ as 0.87 hour at 0.036 nM AuNP in detection of 10 nM sGP. Yet *τ*_*agg*_ could be greatly reduced by increasing AuNP concentration, for example to 0.024 hour, or 36 times shorter, when using 50 times more concentrated NPs.

**Figure 4.**
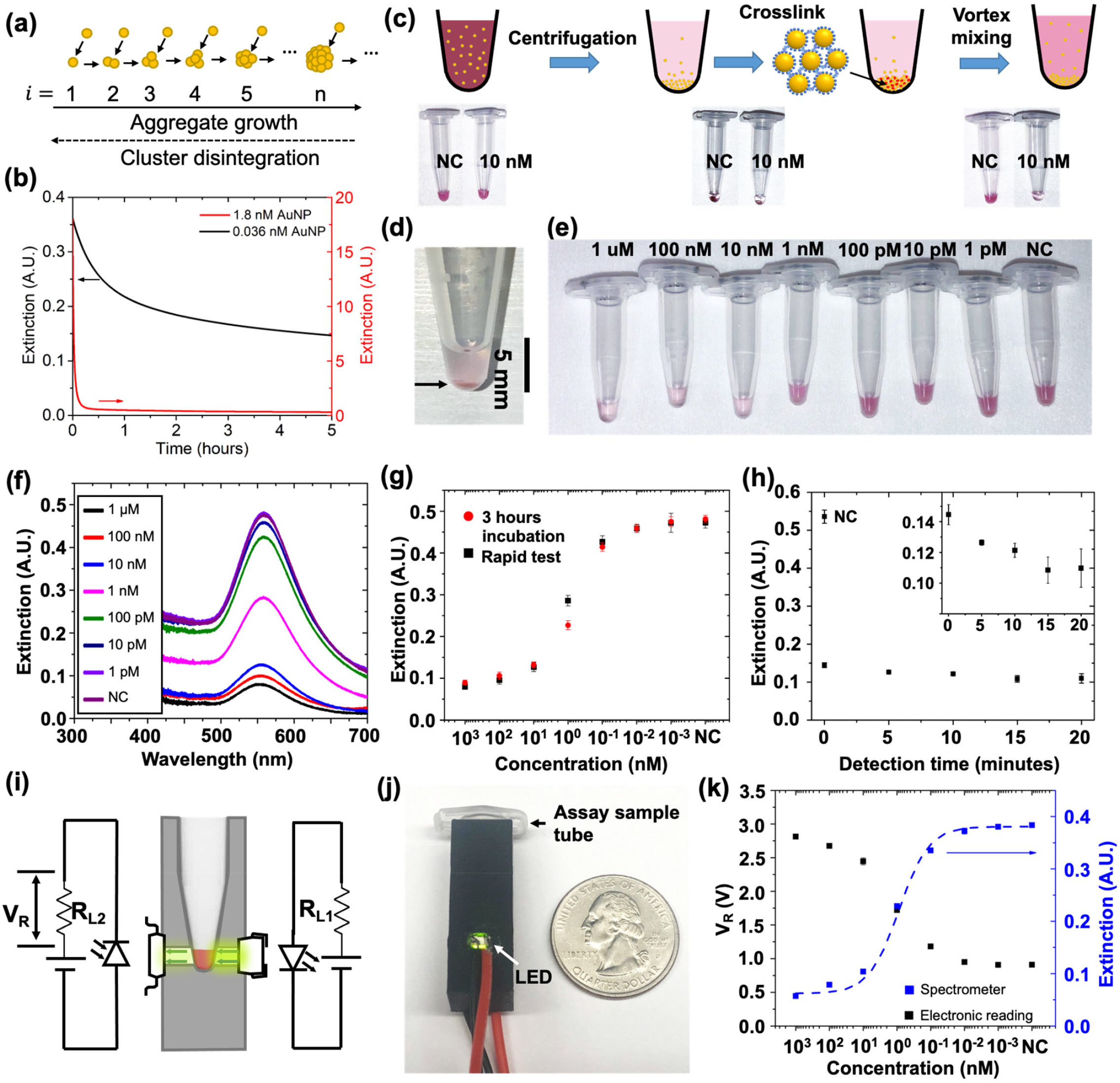
Rapid and electronic detection of sGP using mono-binder antibody sGP49 with improved sensing performance. (a) Modeling of aggregate formation by considering only AuNP monomer-oligomer interactions. (b) Time-dependent extinction calculated for 0.036 nM (black, as used in our experiments) and 1.8 nM (red, 50× concentrated) 80 nm AuNPs in detecting 10 nM sGP. (c) Schematic showing key steps in rapid detection protocol: centrifugation, AuNP aggregation through incubation, and vortex-mixing. (d) Visual image of AuNPs (80 nm, concentration 0.036 nM) after centrifugation at 3,500 rpm for 1 minute. (e) AuNPs in microcentrifuge tubes for rapid detection of 1pM to 1 μM sGP in 5% FBS. The tubes were centrifuged at 3,500 rpm for 1 minute and vortex-mixed for 15 seconds. (f) Extinction spectra of AuNPs shown in (e). (g) Extinction peak values (559 nm) extracted from (f) and plotted against sGP concentration in rapid detection. Measurement results of incubation-based tests were plotted in red for comparison. (h) Effect of incubation time on the extinction measured for samples in detecting 10 nM sGP in 1 × PBS. Inset shows narrowed extinction range for visual contrast. (i-k) Electronic detection of sGP in 5% FBS using miniaturized measurement system: (i) Schematic and (j) Visual image of electronic readout system, consisting mainly of a LED circuit, a photodiode circuit, and a 3D printed Eppendorf tube holder. (k) Voltage signals measured in detecting sGP in 5% FBS, shown as black datapoints. Lab-based spectrometer-measured extinctions of the same assays are plotted in blue.

On the other hand, as gravitational force overcomes fluidic drag, large clusters precipitate to form sedimentation and continuously deplete AuNPs and sGP proteins in the colloid until reaching equilibrium. The sedimentation time can be estimated using the Mason-Weaver equation by *τ*_*sed*_ *z*/(*s* · *g*) where *z* is the precipitation path (for example the height of colloid liquid), *g* is the gravitation constant, and *s* is the sedimentation coefficient dependent on the physical properties of AuNPs and buffers ^38^. Given that *z* ∼3.5 mm for 16 μL liquid in a microcentrifuge tube, we calculated that *τ*_*sed*_ decreases from 26 hours for 80 nm AuNPs to 1.0 and 0.3 hours for a 400 nm and 800 nm diameter cluster (comparable to experimentally observed clusters of micrometer size at 1 nM sGP, Figure S5), respectively. The estimated aggregation and precipitation times are consistent with the experimentally observed incubation time (∼3 hours).

For rapid detection, we introduced a centrifugation step (1,200 × g, 1 min) after antigen mixing to both enhance the reagents’ concentration and decrease the precipitation path (Figure 4**c**, additional data in Supplementary section 7). This step concentrated AuNPs at the bottom of microcentrifugation tubes without causing non-irreversible AuNP aggregation, with an estimated *z* of ∼150 μm as seen from optical image (Figure 4**d**). This corresponds to a roughly >20 times reduction in precipitation path and accordingly *τ*_*sed*_. Additionally, the concentrated AuNPs are confined to an estimated <0.34 μL volume, or ∼50 times concentration increase from original 16 μL colloid liquid, leading to a greatly reduced *τ*_*agg*_, estimated from 0.87 hour to 0.024 hour (Figure 4**b**). These calculations indicate that both the aggregation formation and precipitation of the aggregates can take place in just a few minutes, important to shortening assay time. To experimentally validate the rapid detection concept, the assay colloid was incubated for 20 min after centrifugation and then thoroughly vortexed, which served to re-suspend free AuNPs that could have been physically adsorbed to the tube bottom. Indeed, the increased upper-level assay liquid transparency at higher sGP concentration (Figure 4**e**) was distinguished visually for sGP >1 nM. The extinction values of the upper-level liquid (Figure 4**f**) were extracted at its peak wavelength (∼559 nm) (Figure 4**g**), and plotted against sGP concentration, along with the 3-hour incubation results. The rapid test presented comparable performance in dynamic range and LOD (∼80 pM) compared to incubation. Using 10 nM sGP as the antigen, we found the color contrast was high enough to be immediately resolved by the naked eye after vortex mixing, requiring minimal incubation (Figure 5**h** and S11**a**). Including all of the operation steps for sample collection, pipetting, centrifugation, vortex mixing, and readout, this rapid test scheme can be completed in a few minutes.

**Figure 5.**
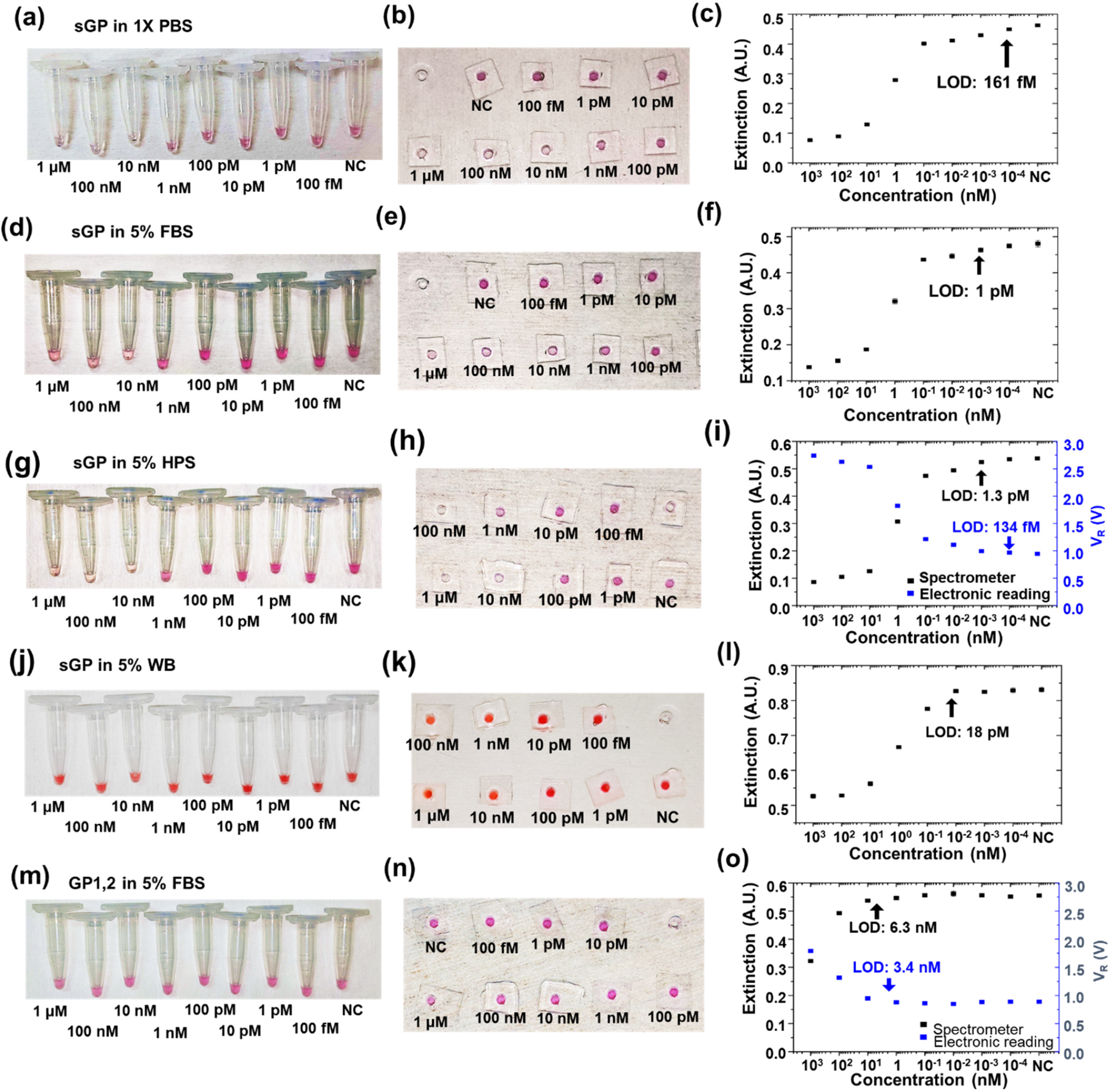
Co-binders sGP49/sGP7 for rapid sGP detection in different buffers. Left column: Optical images of microcentrifuge tubes after rapid test (centrifugation, 20 min incubation, and vortex mixing). Middle column: Optical images of PDMS well plates after rapid test. Right column: Optial sensing (extracted extinction peak values, in black) and electronic sensing (readout voltage, in blue) to detect sGP (or GP1,2) from 100 fM to 1 μM. (a-c) sGP detection in 1× PBS buffer. (d-f). sGP detection in 5% fetal bovine serum (FBS) buffer. (g-i) sGP detection in 5% human pooled serum (HPS) buffer. (j-l) sGP detection in 5% human whole blood (WB). (m-o) GP1,2 detection in 5% FBS buffer.

### Rapid sGP detection with a portable, electronic readout device

Extinction spectrometric analysis provides quantitative and accurate diagnostics but requires bulky spectrometer systems that are more suitable for lab use. We demonstrated the feasibility of detecting sGP in FBS using a cost-efficient, portable UV-visible spectrometer system for field deployment (Figure S13). Additionally, we developed a homemade LED-photodiode based electronic readout system with significantly reduced system cost to deliver accurate and sensitive detection results comparable to a lab-based spectrometer system (Figure 4 **i-k**). Here, an LED light source emitted narrowband light at the AuNP extinction peak (*λ*_*P*_ 560 nm, *FWHM*_*P*_ = 40 nm), which transmitted through the upper-level assay liquid and then was collected by a photodiode (Figure 4**i**). As a result, a photocurrent or photovoltage was generated on a serially connected load resistor in a simple circuitry that can be easily integrated and scalably produced. In practice, we 3D-printed a black holder to snug-fit a microcentrifuge tube, and mounted the LED and photodetector on two sides of the holder (Figure 4**j**). The LED and photodiode were powered by alkaline batteries (3V and 4.5V, respectively), and the bias voltages were set to ensure wide-range detection of sGP proteins without saturating the photodetectors. Using 80 nm sGP49-functionalized AuNP Nano2RED assays, sGP was detected in diluted FBS (5%) by reading the photovoltage signals with a handheld multimeter (Figure 5**k**, *V*_*R*_, in black). Compared to lab-based spectrometric readout (in blue), the electronic readout displayed identical dynamic range, but slightly improved LOD (27 pM compared to 80 pM).

### Detection of sGP and RBD in serum and blood

We further evaluated the use of co-binding nanobodies in sGP and RBD sensing (Figures 5 and 6), *i*.*e*., sGP49/sGP7 for sGP and RBD8/RBD10 for RBD (Figure 2), and performed Nano2RED tests in different buffers, including PBS, FBS, human pooled serum (HPS), and whole blood (WB) (additional data in Supplementary section 8 and Figures S15 to S23). The incubation and rapid assay formats with the assay performance, instrument costs, and LODs were summarized in Table1 and 2, and additional data on measurement variance (sigma) and LOD were summarized in Supplementary Tables S1 and S2. There are several notable observations. First, using sGP sensing as an example and comparing to previously reported results ^18, 39^ (Supplementary Table S2), Nano2RED with spectrometric and electronic readout consistently produced ∼130 fM to 1.3 pM LOD (or ∼10 to ∼100 pg/mL) in PBS, FBS, and HPS. It is noted that a very recently reported co-binder-based D4 assay format reported ∼30 pg/mL LOD in human serum, and was able to detect the Ebola virus earlier than PCR in a monkey mode□^18^. In LOD comparison, the sensitivity of Nano2RED (∼10 pg/mL with electronic readout) is even better, indicating its competitiveness in high-precision diagnostics.

**Figure 6.**
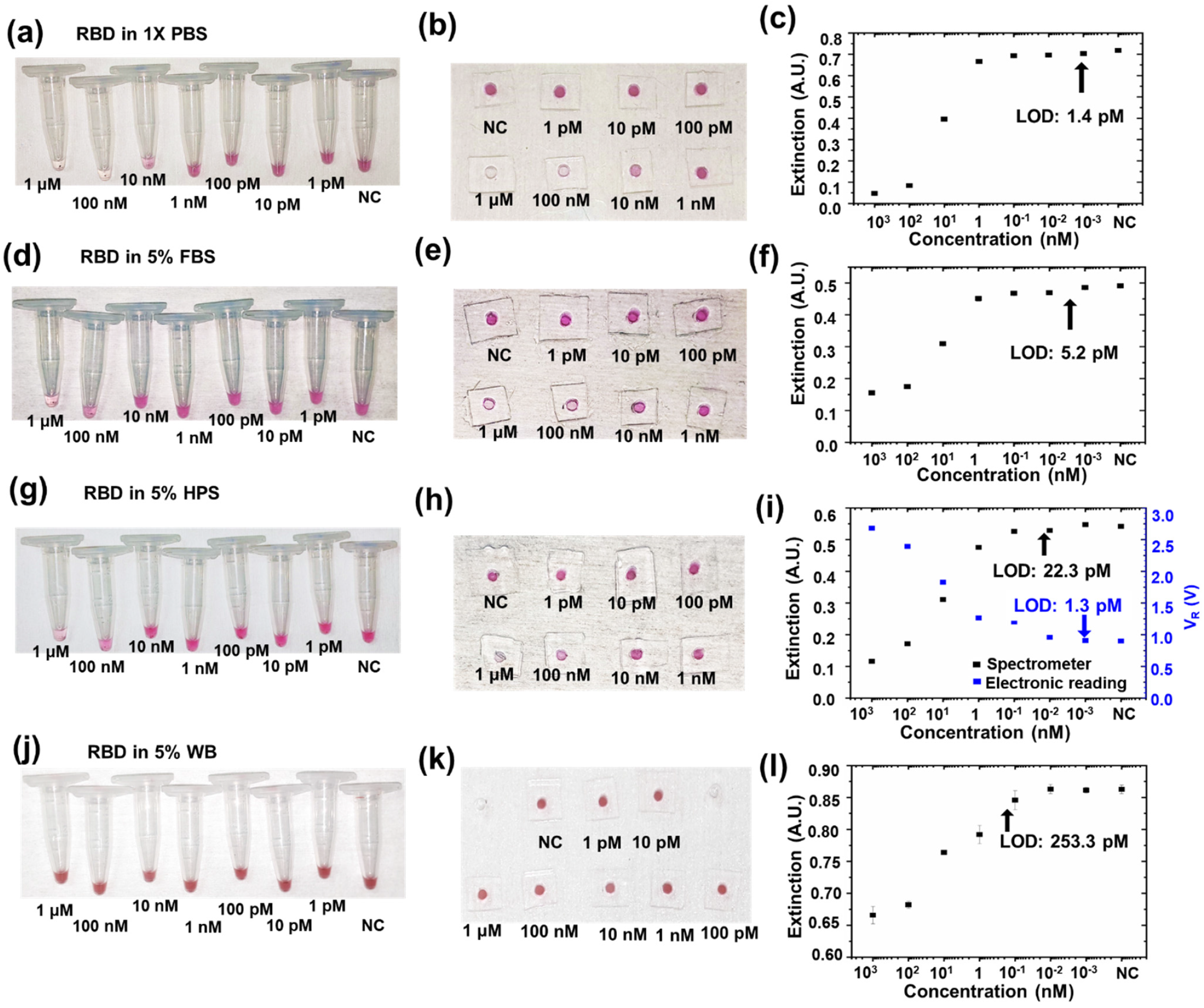
Co-binders R8/R10 for rapid RBD detection in different buffers. Left column: Optical images of microcentrifuge tubes after rapid test (centrifugation, 20 min incubation, and vortex mixing). Middle column: Optical images of PDMS well plates after rapid test. Right column: Optial sensing (extracted extinction peak values, in black) and electronic sensing (readout voltage, in blue) to detect RBD from 1 pM to 1 μM. (a-c) RBD detection in 1× PBS buffer. (d-f). RBD detection in 5% FBS buffer. (g-i) RBD detection in 5% HPS buffer. (j-l) RBD detection in 5% WB buffer.

**Table 2.** Performance of Nano2RED in SARS-CoV-2 RBD protein sensing.

| Assay format | Readout method | Readout system | Readout system cost | Readout system size | Assay time | Buffer | Diagnostic sensitivity |  |
| --- | --- | --- | --- | --- | --- | --- | --- | --- |
|  |  |  |  |  |  |  | pM | pg/mL |
| Centrifuge-accelerated (rapid) | Colorimetric | Eyes or smart phones | ~\$200 (phone) | ~78 cm<sup>3</sup> | 5-20 min | FBS | 1000 | 29180 |
| | Spectrometric | Lab microscope | \$25,000 | ~3 m<sup>3</sup> | | PBS | 1.4 | 42 |
|  |  |  |  |  |  | FBS | 5.23 | 153 |
|  |  |  |  |  |  | HPS | 22.3 | 649 |
|  |  |  |  |  |  | WB | 253.3 | 7372 |
| | Electronic | LED & photodetector | ~\$5 | 1.1 × 1.1 × 3.3 cm<sup>3</sup> | | HPS | 1.35 | 39 |

Additionally, our study (Table 1) also revealed the importance of a systematic assay design strategy, from molecular binding to signal transduction and readout, to optimize antigen detection. It is clear that the co-binder pair improved the LOD by 10 to 100 times compared to the mono-binder (sGP49, *k*_*D*_ 4.6 nM), despite a relatively low *k*_*D*_ of 199 nM for the second binder (sGP7) (Figure 2). This improved sensitivity is likely because the co-binders have a favorable, non-competitive binding configuration that serves to improve antigen binding and AuNP aggregation. Uniquely, the use of a portable and inexpensive electronic readout did not negatively affect the LOD of Nano2RED, but rather improved it compared to spectrometric readout (Figures 5**i** and 6**i**, and Table 1 and 2). This can be attributed to smaller 3-sigma errors in the electronic readout (Supplementary Table S2), partly due to a larger signal fluctuation in optical imaging caused by manual operation, such as in focusing. Here, the electronic signal is mainly dependent on the circuit elements but much less dependent on operators’ judgement, and thus potentially more reliable and accurate. In addition, the use of biological buffers could also affect detection. For both Ebola sGP and SARS-CoV-2 RBD, the LOD increased by about 5-10 times in serum (FBS and HPS) than in PBS, and further increased by another 10 times in WB. Additionally, the colorimetric readout by the naked eye was capable of detecting both antigens in serum at concentrations higher than 100 pM or 1 nM (Figures 5 and 6); however, it became challenging to do so in WB, mainly due to the fact that WB absorbs in short wavelengths and causes background color interference with AuNPs (Figures 5 **j-k**). However, the spectrometric readout could still readily identify the sGP or RBD extinction signals from the background WB absorption for accurate detection (Figure S19**e** and Figure 5**l** for sGP and Figure S25 **e** and Figure 6**l** for RBD), indicating the feasibility of Nano2RED for field use with minimized sample preparation.

Fundamentally different from conventional high-sensitivity antigen diagnostics that usually require bulky and expensive readout systems, as well as long assay time, Nano2RED is an affordable and accessible diagnostic technology. For example, ultrasensitive sGP sensing using NP-enhanced fluorescent readout would require 3-4 hours of image processing to reduce noise for optimal sensing^39^, and these fluorescent systems usually require cubic meter space and costb$40,000 or more (a high-end fluorescent camera with high signal-to-noise ratio is ∼$25,000). Similarly, a D4 co-binder assay requires a lab-based bulky fluorescent system and ∼60 min assay time to achieve PCR-comparable diagnostic sensitivity. Its sensing performance drops ∼10 times to 100 pg/mL when using a customized fluorescent system, which costs ∼$1,000 and occupies ∼3,000 cm^3^. The performance further decreases to 6,000 pg/mL when using LFA with colorimetric readout^18^. Clearly standing apart from the rest, Nano2RED utilizes miniaturized and low-cost semiconductor devices for signal readout rather than a fluorescent system. Therefore, it has a very small footprint (4 cm^3^ for tube holder, or <100 cm^3^ for the whole system, including batteries and meters, which all could be miniaturized on a compact circuit board in the future), is very low cost (LED and photodiode each <$1 here, but can be <$0.1 when used at large scale, with the total system cost estimated well below $5), and offers a rapid readout (5 to 20 min, depending on incubation time after centrifugation). Further, the electronic readout is more accurate than the colorimetric readout, more accessible, without color vision limitations, and more readily available for data storage in computers or online databases for real-time or retrospective data analysis. Additionally, we have estimated the reagent cost in Nano2RED is only about $0.01 per test (Supplementary section 9), since it requires only a small volume (∼20 μL) of regents.

We tested sGP against GP1,2, a homotrimer glycoprotein transcribed from the same GP gene and sharing its first 295 residues with sGP ^16^, both in FBS (Figures 5 **d-f** and **m-o**). The majority of GP1,2 can be found on virus membranes whereas a small portion is released into the patient’s bloodstream. The close relevance of GP1,2 to sGP makes it a very strong control molecule to assess our assay’s specificity. Indeed, GP1,2 did not produce detectable signals unless higher than 1 nM, indicating a high selectivity over a broad concentration range (100 fM to 1 nM, or 4 logs) where minimal nanobody binding or AuNP aggregation occurred. A high assay specificity is crucial for minimizing false positive diagnosis of infectious diseases, which could lead to unnecessary hospitalizations and even infections. Considering that 10 nM and higher sGP concentration is typical for EVD patients^32, 33, 40^, Nano2RED is particularly suitable for high-speed mass screening of EVD susceptible populations. Further, SARS-CoV-2 RBD proteins were also detected in the single-digit picomolar range in PBS, FBS, and HPS, with the best LOD (Table 2, 1.3 pM, or ∼40 pg/mL) again achieved with electronic readout. The LOD in RBD sensing is ∼10 times higher compared to sGP sensing, mainly attributed to lower binding affinities of the nanobodies obtained from a single-round biopanning (Figure 2). Tighter binders can be selected using more biopanning rounds; however, the detection of RBD, a monomeric protein target, serves to demonstrate the general feasibility of the Nano2RED co-binding assay in detection of a broad range of antigens, regardless of their complex molecular structures. Considering the fact that the spike protein is a trimer and each SARS-CoV-2 particle is covered with ∼20 copies of such trimers^41^, the detection of SARS-CoV-2 virus particles could behave differently. The detection of SARS-CoV-2 particles might have a better sensitivity. The detection of viral particles from patient samples would itself be quite exciting future studies but beyond the scope of this work. Nevertheless, the broadband dynamic range (∼100 fM to 1 μM for co-binders, or 7 logs), high sensitivity, high specificity, and broad applications therefore make our Nano2RED assay highly feasible for precise antigen quantification and detection of early-stage infection.

### Conclusions and Outlook

We have demonstrated a generalizable and rapid assay design and pipeline that combines fast affinity reagent selection and production with nanometer-scale theoretical analysis and experimental characterization for optimized sensing performance. Synthetic, high-affinity, co-binding nanobodies, which could be quickly produced by a phage display selection method from a premade combinatorial library for any given antigen, proved to be effective in detecting dimeric Ebola sGP and monomeric SARS-CoV-2 RBD proteins. The Nano2RED utilized unique signal transduction pathways to convert biological binding into electronic readout. Using simple electronic circuitry, it starts with AuNP aggregation (governed by dynamic antigen-nanobody interactions described by Langmuir isotherm), which triggers AuNP cluster sedimentation (explained by Mason-Weaver equation^42^), and then enables AuNP-concentration-dependent optical extinction (following Beer-Lambert law). The use of AuNPs for in-solution testing serves to greatly facilitate fluidic transport and antigen binding at nanometer scale. Nano2RED eliminates the need for long-time incubation due to slow analyte diffusion in conventional ELISA and other plane surface-based assays, as well as its associated cumbersome washing steps. The introduction of brief centrifugation and vortex mixing further greatly shortens the aggregation and sedimentation time, enabling rapid tests (within 5 to 20 min) without sacrificing sensitivity or specificity. Our data showed that Nano2RED is highly sensitive (sub-picomolar or ∼10 pg/mL level for sGP) and specific in biological buffers while also affordable and accessible. Importantly, the portable electronic readout, despite being very simple and inexpensive (<$5), proved to be more reliable and sensitive than colorimetric and even spectrometric readout. Nano2RED can be applied for lab tests to detect early-stage virus infection at a high sensitivity potentially comparable to PCR. It can also be used for high-frequency at-home or in-clinic diagnostics, as well as in resource-limited regions, which could greatly enhance control of disease transmission. The digital data format will also reduce human intervention in data compiling and reporting, while facilitating fast and accessible data analysis. Nano2RED may find immediate use in the current COVID-19 pandemic for both antigen and antibody detection, as well as preparing for future unforeseeable new outbreaks.

## METHODS

### Materials

Phosphate-buffered saline (PBS) was purchased from Fisher Scientific. Bovine serum albumin (BSA) and molecular biology grade glycerol were purchased from Sigma-Aldrich. Fetal bovine serum (FBS) was purchased from Gibco, Fisher Scientific. FBS was used without heat inactivation to best reflect the state of serum collected in field. Polyvinyl alcohol (PVA, M_W_ 9,000-10,000) was purchased from Sigma-Aldrich. Sylgard 184 silicone elastomer kit was purchased from Dow Chemical. DNase/RNase-free distilled water used in experiments was purchased from Fisher Scientific. Phosphate Buffered Saline with Tween 20 (PBST), Nunc MaxiSorp 96 well ELISA plate, streptavidin, 1% casein, 1-Step Ultra TMB ELISA substrate solution, and isopropyl-β-D-galactopyranoside (IPTG) were purchased from Thermo Fisher Scientific. HRP-M13 major coat protein antibody was purchased from Santa Cruz Biotechnology. Sucrose and imidazole were purchased from Sigma-Aldrich. A 5 mL HisTrap column, HiLoad 16/600 Superdex 200 pg column, and HiPrep 26/10 desalting column were purchased from GE Healthcare. BirA-500 kit was purchased from Avidity. Streptavidin (SA) Biosensors were purchased from ForteBio. The streptavidin functionalized AuNPs, dispersed in 20% v/v glycerol and 1 wt% BSA buffer, were purchased from Cytodignostics. Thiolated carboxyl polyethylene glycol linker was self-assembled on AuNP through a thiol-sulfide reaction. Streptavidin was then surface functioned through amine-carboxyl coupling by N-Hydroxysuccinimide/1-Ethyl-3-(3-dimethylaminopropyl) carbodiimide (NHS/EDC) chemistry.

### Phage display selection

sGP and RBD (GenScript) protein binder selection was done according to previously established protocols^15^. In brief, screening was performed using biotin and biotinylated target protein-bound streptavidin magnetic beads for negative and positive selections, respectively. Prior to each round, the phage-displayed nanobody library was incubated with the biotin-bound beads for 1 h at room temperature to remove off-target binders. Subsequently, the supernatant was collected and incubated with biotinylated-target protein-bound beads for 1 h. Beads were washed with 10× 0.05 % PBST (1×PBS with 0.05% v/v Tween 20) and phage particles were eluted with 100 mM triethylamine. A total of three rounds of biopanning were performed with decreasing amounts of antigen (200 nM, 100 nM, 20 nM). Single colonies were picked and validated by phage ELISA followed by DNA sequencing.

### Single phage ELISA

ELISAs were performed according to standard protocols^15^. Briefly, 96 well ELISA plates (Nunc MaxiSorp, Thermo Fisher Scientific) were coated with 100 μL 5 μg/mL streptavidin in coating buffer (100 mM carbonate buffer, pH 8.6) at 4°C overnight. After washing with 3× 0.05 % PBST (1×PBS with 0.05% v/v Tween 20), each well was added to 100 μL 200 nM biotinlyated target protein and incubated at room temperature for 1 h. Each well was washed by 5× 0.05 % PBST, blocked by 1% casein in 1×PBS, and added to 100 μL single phage supernatants. After 1 h, wells were washed by 10×0.05% PBST, added to 100 μL HRP-M13 major coat protein antibody (RL-ph1, Santa Cruz Biotechnology; 1:10,000 dilution with 1×PBS with 1% casein), and incubated at room temperature for 1 h. A colorimetric detection was performed using a 1-Step Ultra TMB ELISA substrate solution (Thermo Fisher Scientific) and OD450 was measured with a SpectraMax Plus 384 microplate reader (Molecular Devices).

### Mono-binders purification and biotinlyation

sGP7, sGP49, RBD8, and RBD10 mono-binders were expressed as a C-terminal Avi-tagged and His-tagged form in *E. coli* and purified by Ni-affinity and size-exclusion chromatography. In brief, *E. coli* strain WK6 was transformed and grown in TB medium at 37°C to an OD600 of ∼0.7, then induced with 1 mM isopropyl-β-D-galactopyranoside (IPTG) at 28°C for overnight. Cell pellets were resuspended in 15 mL ice-cold TES buffer (0.2 M Tris-HCl pH 8.0, 0.5 mM EDTA, 0.5 M sucrose) and incubated with gently shaking on ice for 1 h, then added to 30 mL of TES/4 buffer (1:4 dilution of the TES buffer in ddH_2_O) and gently shaken on ice for 45 min. Cell debris was removed by centrifugation at 15,000×g, 4°C for 30 mins. The supernatant was loaded onto a 5 mL HisTrap column (GE Healthcare) pre-equilibrated with the lysis buffer (50 mM sodium phosphate, pH 8.0, 300 mM NaCl, 10 mM imidazole, 10% glycerol). The column was washed with a washing buffer (50 mM sodium phosphate, pH 8.0, 300 mM NaCl, 20 mM imidazole, 10% glycerol) and then His-tagged proteins were eluted with an elution buffer (50 mM sodium phosphate, pH 8.0, 300 mM NaCl, 250 mM imidazole, 10% glycerol). Eluates were loaded onto a HiLoad 16/600 Superdex 200 pg column (GE Healthcare) pre-equilibrated with a storage buffer (1×PBS, 5% glycerol). Eluted proteins were concentrated, examined by SDS-PAGE, and quantified by a Bradford assay (BioRad), then flash frozen in 100 μL aliquots by liquid N_2_ and stored at -80°C.

The purified protein was biotinylated by BirA using a BirA-500 kit (Avidity). Typically, 100 μL BiomixA, 100 μL BiomixB, and 4 μL 1 mg/mL BirA were added to 500 μL protein (∼1 mg/mL), and adjusted to a final volume of 1000 μL with nuclease-free water (Ambion). The biotinylation mixture was incubated at room temperature for 1 h and then loaded onto a HiPrep 26/10 desalting column (GE Healthcare) pre-equilibrated with a storage buffer (1×PBS, 5% glycerol) to remove the free biotin.

### Binding kinetics analysis

The four mono-binders’ binding kinetics were analyzed using an Octet RED96 system (ForteBio) and Streptavidin (SA) biosensors. 200 nM biotinylated sGP or RBD target protein was immobilized on SA biosensors with a binding assay buffer (1×PBS, pH 7.4, 0.05% Tween 20, 0.2% BSA). Serial dilutions of mono-binder were used for the binding assay. Dissociation constants (*k*_*D*_) and kinetic parameters (*k*_*on*_ and *k*_*off*_) were calculated based on global fit using Octet data analysis software 9.0. For co-binder validation, sGP or RBD bound SA biosensors were first dipped into the sGP49 or RBD10 wells 750 s for saturation, then incubated with sGP7 or RBD8 for 750 s.

### Determination of detection sensitivity using a co-binder sandwich ELISA assay

In order to validate and determine the detection sensitivity of co-binders, a sandwich ELISA-like assay was performed. Briefly, 96 well ELISA plates (Nunc MaxiSorp, Thermo Fisher Scientific) were coated with 100 μL 5 μg/mL streptavidin in coating buffer (100 mM carbonate buffer, pH 8.6) at 4°C overnight. After washing with 3× 0.05 % PBST (1×PBS with 0.05% v/v Tween 20), each well was added to 100 μL 200 nM biotinlyated sGP49 or RBD10 (∼100 nM) protein and incubated at room temperature for 1 h, then washed with 5× 0.05 % PBST and added to 100 μL serial dilutions (0 to 500 nM) of sGP or RBD protein and incubated at room temperature for 1 h. Each well was subsequently blocked by 1% casein in 1×PBS for 1 h, then added to 100 μL sGP7 or RBD8 phage supernatants. After 1 h, wells were washed by 10× 0.05% PBST, added to 100 μL HRP-M13 major coat protein antibody (RL-ph1, Santa Cruz Biotechnology; 1:10,000 dilution with 1×PBS with 1% casein), and incubated at room temperature for 1 h. A colorimetric detection was performed using a 1-Step Ultra TMB ELISA substrate solution (Thermo Fisher Scientific) and OD450 was measured with a SpectraMax Plus 384 microplate reader (Molecular Devices). Limit of Detection (LoD) was calculated by mean blank + 3×SD_PS4_. More details on the SD selection is provided in supplementary section (Tables S1 and S2, and Supplementary Section 8.4 and 8.5)

### Nanoparticle functionalization with nanobodies

The streptavidin surface functioned gold nanoparticles (typically ∼0.13 nM 80 nm AuNPs, 80 μL) were first mixed with an excessive amount of biotinylated nanobodies (about 1.2 μM, 25 μL). The mixture was then incubated for 2 hours to ensure complete streptavidin-biotin conjugation. Next, the mixture was purified by centrifuge (accuSpin Micro 17, Thermo Fisher) at 10,000 rpm for 10 min and repeated twice to remove unbounded biotinylated nanobodies. The purified AuNP colloid was measured by Nanodrop 2000 (Thermo Fisher) to determine the concentration. The concentration of AuNP in colloid was subsequently adjusted to get an optimal extinction level (*e*.*g*., empirically 0.048 nM for 80 nm AuNPs) and was aliquoted into 12 uL in a 500 uL Eppendorf tube.

### Antigen detection

Target sGP or RBD stock solution (6 μM, in 1×PBS) underwent a 10-fold serial dilution to target concentrations (4 pM to 4 uM) in selected detection media. The final composition of PBS detection media was composed of 1×PBS, 20% v/v glycerol and 1 wt% BSA while that of FBS (Fetal Bovine Serum), HPS (Human Pooled Serum), and WB (Whole blood) detection media had a final concentration of 1×PBS, 20% v/v glycerol and 1 wt% BSA and 20% of either FBS, HPS, or WB, which resulted in a final concentration of FBS, HPS, or WB in the detection assay to be 5%. For example, for sGP detection with co-binders, solutions of sGP49-functionalized AuNPs, sGP7-functionalized AuNP, and sGP were mixed in a 500 uL Eppendorf tube at a ratio of 3:3:2 and thoroughly vortexed. When incubation-based detection was used, the solution was allowed to incubate (typically 3 hours) prior to readout. A similar protocol was used for RBD sensing.

### Centrifuge enhanced rapid detection

The same protocols were followed in preparation of sGP49 surface functioned AuNP colloid. After mixing sGP49 surface functioned AuNP colloid with sGP solution, the AuNP assay colloid was centrifuged (accuSpin Micro 17, Thermo Fisher) at 3,500 rpm (1,200×g) for 1 minute. After optional incubation, the assay colloid was vortexed (mini vortexer, Thermo Fisher) at 800 rpm for 15 seconds prior to readout. A similar protocol was used for RBD sensing.

### PDMS well plate fabrication

Sylgard 184 silicone elastomer base (consisting of dimethyl vinyl-terminated dimethyl siloxane, dimethyl vinylated, and trimethylated silica) was thoroughly mixed with the curing agent (mass ratio 10:1) for 30 minutes and placed in a vacuum container for 2 hours to remove the generated bubbles. The mixture was then poured into a flat plastic container at room temperature and incubated for one week, until the PDMS is fully cured. The PDMS membrane was then cut to rectangular shape, and 2 mm wells were drilled by punchers. To prevent non-specific bonding of proteins, the PDMS membrane was treated with PVA, adapted from methods described by Trantidou et al.^43^ The as-prepared PDMS membrane and a diced rectangle shaped fused silica (500 μm thick) were both rinsed with isopropyl alcohol, dried in nitrogen, and treated by oxygen plasma (flow rate 2 sccm, power 75 W, 5 min). Immediately after, the two were bonded to form a PDMS well plate. The plate was further oxygen plasma treated for 5 min and immediately soaked in 1% wt. PVA in water solution for 10 min. Then, it was dried by nitrogen, heated on a 110 °C hotplate for 15 min, and cooled to room temperature by nitrogen blow.

### UV-visible spectrometric measurement and dark field scattering characterizations

The UV-visible spectra and dark field imaging were performed using a customized optical system (Horiba), comprising an upright fluorescence microscope (Olympus BX53), a broadband 75W Xenon lamp (PowerArc), an imaging spectrometer system (Horiba iHR320, spectral resolution 0.15 nm), a low-noise CCD spectrometer (Horiba Syncerity), a high-speed and low-noise EMCCD camera (Andor iXon DU897 Ultra), a vision camera, a variety of filter cubes, operation software, and a high-power computer. For spectral measurement, the PDMS plate loaded with upper-level assay samples or drop-cast samples on a glass slide was placed on the microscope sample stage. Light transmitted through PDMS well plate was collected by a 50×objective lens (NA=0.8). The focal plane was chosen at the well plate surface to display the best contrast at the hole edge. A 10×objective lens (NA=0.3) was used for drop-cast samples. The signals were typically collected from the 350 nm to 800 nm spectral range with integration time of 0.01 s and averaged 64 times. The dark field scattering was illuminated by an xeon lamp, collected by a 100×dark field lens (NA=0.9), and imaged by an EMCCD camera. The integration time was set to 50 ms. For each drop-cast sample spot, ten images were taken from different areas in the spot. The size for the area taken in each dark-field scattering image was 62.5 μm×62.5 μm.

### SEM imaging of drop-cast samples on gold

The SEM image was taken by a Hitachi S4700 field emission scanning electron microscope at 5 kV acceleration voltage and magnification of 15,000. For each drop-cast sample spot, ten images were taken from different regions in the spot. The size of the region taken in each SEM image was 8.446 μm×5.913 μm.

### TEM to image AuNP precipitates

The upper-level assay liquid was removed from the microcentrifuge tube until 2 to 3 μL of sample containing AuNP precipitates were left. The tube was vortexed thoroughly, and 2 μL of remaining samples were pipetted, and coated on a Cu grid (Electron Microscopy Sciences, C flat, hole size 1.2 μm, hole spacing 1.3 μm) that was pre-treated on both sides with oxygen plasma (30 seconds). The Cu grid was plunge frozen in ethane using Vitrobot plunge freezer (FEI). The blot time was set to 6 sec. After plunging, the sample was soaked in liquid nitrogen for long-term storage. FEI Tecnai F20 transmission electron microscope (200 kV accelerating voltage) was used for CryoTEM imaging. 25 high-resolution TEM images were taken for 1 μM, 1 nM sGP in PBS samples and reference samples, respectively. The size of the area taken in each image was 4.476 μm×4.476 μm.

### Portable spectrometric readout

The assay colloids were initially characterized by a miniaturized portable UV-visible spectra measurement system. OSL2 fiber coupled illuminator (Thorlabs) was used as the light source. The light passed through the 4 mm diameter wells loaded with assay colloid and coupled to Flame UV-visible miniaturized spectrometer (Ocean optics) for extinction spectra measurement. The signals were averaged from six scans (each from 430 nm to 1100 nm) and integrated for 5 seconds in each scan.

### Electronic readout with rapid test

A LED-photodiode electronic readout system consists of three key components: a LED light source, a photodiode, and a microcentrifuge tube holder. The centrifuge tube holder was 3D printed using ABSplus P430 thermoplastic. An 8.6 mm diameter recess was designed to snuggly fit a standard 0.5 mL Eppendorf tube. 2.8 mm diameter holes were open on two sides of the microcentrifuge tube holder to align a LED (597-3311-407NF, Dialight), the upper-level assay liquid, and a photodiode (SFH 2270R, Osram Opto Semiconductors). The LED was powered by two Duracell optimum AA batteries (3 V) through a serially connected 35 Ω resistor to set the LED operating point. The photodiode was reversely biased by three Duracell optimum AA batteries (4.5 V) and serially connected to a 7 MΩ load resistor. The photocurrent that responds to intensity of light transmitted through the assay was converted to voltage through the 7 MΩ load resistor and measured with a portable multimeter (AstroAI AM33D).

### Estimate of limit of detection for Nano2RED

In our work, limit of detection (LOD) was calculated according to the International Union of Pure and Applied Chemistry definition, that is, the concentration at which the measured response is able to distinguish from the reference signal by three times the standard deviation in measurements. For optical measurement, we used *LoD* = *c*(*E*_*NC*_ − 3*σ*). Here, the reference was averaged over three measurements of the negative control (NC) sample. Whereas for electronic measurement, we used *LoD c*(*V*_*NC*_ + 3*σ*), where *V*_*NC*_ is the readout voltage for NC sample. We compared different methods to estimate *σ, i*.*e*., the conventional way considering only the NC sample (*σ*_*NC*_), using pooled sigma from all measurements (*σ*_*PSA*_), and pooled sigma from the four lowest concentrations (*σ*_*PS4*_) (Table S1 and Figure S25). We noticed in our case that using *σ*_*PSA*_ and *σ*_*PS4*_ both yielded more consistent reporting of LOD compared to using *σ*_*NC*_, and therefore the sensor LOD was estimated with *σ*_*PSA*_ for its best consistency (Table 1). This consistency could be attributed to the nature of our data collection is a physical process and less dependent on reagent concentration compared to conventional ELISA. Particularly when using spectrometric readout, the noise is strongly affected by optical focusing and could happen to any data sets; and therefore the overall average provides a better estimate of the empirical errors. Differently, for ELISA measurement, the sigma is much smaller at lower concentrations, so we used conventional *σ*_*NC*_ for LOD determination.

## Acknowledgements

This work was supported by a grant from the U.S. National Institutes of Health (1R35GM128918) to L. Gu., C. Wang, X. and Chen, M.A. Ikbal and J. Zuo acknowledge partial support from National Science Foundation under grant no. 1809997, 1847324, and 2020464. The samples were characterized in the Eyring Materials Center (EMC) at Arizona State University. Access to the EMC was supported, in part, by NSF grant no. 1542160. We thank Y. Yao for the use of the portable spectrometer system, B. Ipema and S. Myhajlenko for support in electronic measurement setup, D. Baker and L. Stewart for providing recombinant sGP and GP1,2 proteins, Y. Liang for the help on sGP biopanning, and D. Williams for cryoTEM inspections.

## Author contributions

X. C., S. K., and M. A. I. contributed equally. X.C. performed all the experiments for incubation-based Ebola sGP sensing, characterized the sensing performance by microscopy and spectroscopy, modelled the sensing mechanism, established rapid detection and electronic readout process, and contributed to manuscript writing. M. A. I. conducted sGP and RBD detection using co-binders in biological buffers and analyzed the sensing performance. S. K. performed sGP and RBD co-binders screening. S. K. and Y. P. biochemically characterized co-binder candidates. Z. Z. performed initial sensing experiments to study spectroscopic readout in sGP detection. J. Z. contributed to the portable spectroscopic setup. L.G. conceived the idea of using synthetic nanobodies for antigen (sGP and RBD) detection and led the efforts in screening and characterization of nanobodies. C. W. conceived the idea of using NP for in-solution antigen assay, investigated the sensing mechanism, designed the incubation-based and rapid assay formats, led experiments to study the sGP and RBD sensing performance with colorimetric, spectrometric, and electronic readout. C.W. and L.G. coordinated this research project and wrote the manuscript together. Correspondence and requests for materials should be addressed to L.G. or C.W.

## Supplementary information

The online version contains supplementary material available at XXX.

## Notes

### Competing Interest Statement

The authors have filed patent applications related to the publication

### Summary of Updates

We have performed additional tests using cobinding nanobodies towards Ebola sGP protein sensing, which showed greatly improved sensitivity. We also added data for testing of SARS-CoV-2 RBD protein using cobinding nanobodies. The manuscript has been edited throughput.

