## Supplementary document for "Synthetic Nanobody-Functionalized Nanoparticles for Accelerated Development of Rapid, Accessible Detection of Viral Antigens"

### 1. Characterization of nanobody binders

(a)

| Co-binder ID | CDR1 | CDR2 | CDR3 |
|---|---|---|---|
| sGP7 | FSSSNYA | RWDNVKA | MVDDYWDPG |
| sGP49 | DTYYIYA | RSSSAAD | SWHVRFDDYEKVISMQEV |
| RBD8 | SYSEFES | WLWGNQV | EKHWMQYDF |
| RBD10 | SGSFASV | FFENTIT | YWMVVQHNMSPD |

| Co-binder ID | $K_D$ ($10^{-9}$ M) | $k_{on}$ ($10^3 M^{-1}s^{-1}$) | $k_{off}$ ($10^{-4}s^{-1}$) |
|---|---|---|---|
| sGP7 | 199 | 2.97 | 5.91 |
| sGP49 | 4.63 | 40.7 | 1.88 |
| RBD8 | 29.8 | 286 | 85.4 |
| RBD10 | 613 | 1.50 | 9.22 |

(c)

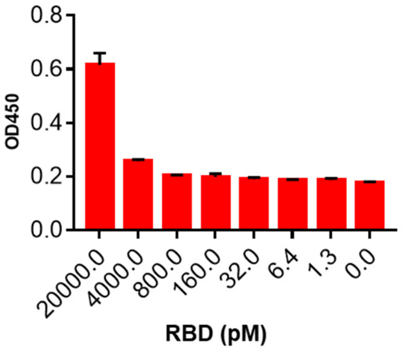


(b)

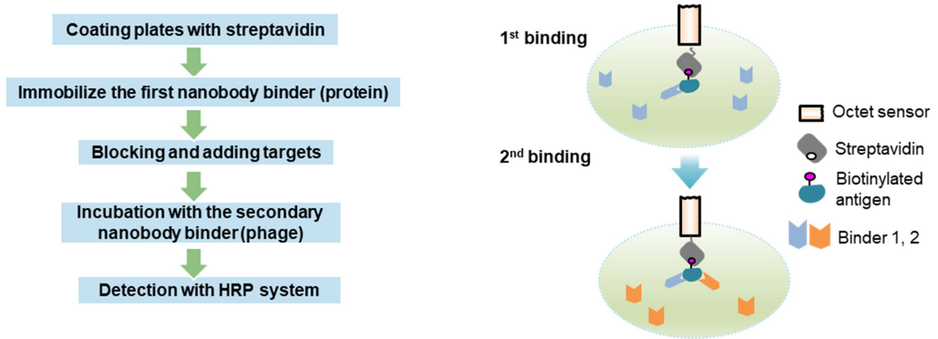


**Figure S1. Characterization of co-binders for sGP and RBD antigen detection.** (**a**) CDR sequences and dissociation constants of top two co-binders identified in this work. (**b**) Schematic of the co-binder validation methods: ELISA (left panel) and BLI (right panel). (**c**) The detection of RBD spiked in PBS buffer with co-binders. Limits of detection (LoD) is given in more details in supplementary section 8.4.

### 2. Finite-difference time-domain (FDTD) simulation of AuNP extinction

We performed FDTD simulation of different numbers of AuNPs in a cluster (Figure S2). AuNP cluster with given number gold nanoparticles were modeled using densely packed 80 nm AuNP nanoparticles with a spacing of 12.8 nm (sGP49-sGP-sGP49 bridge length). The boundary condition along *x*-, *y*- and *z*- direction was set as perfect matched layer (PML). A total-field scattered-field (TFSF) light source (400-1000 nm) was used to calculate extinction cross section. The mesh size was set to be 5 nm in *x*-, *y*- and *z*- direction in the AuNP cluster region. Six monitors recording the power flux were set outside TFSF source region normal to *x*-, *y*- and *z*- directions. The background index was set as 1.33 to simulate the solution environment. The simulation time was set to 5000 fs and auto shut off threshold was set as 5 $\times 10^6$.

The extinction cross sections per AuNP for AuNP clusters with different AuNP number is plotted in Figure S2. We found the resonance peaks red-shift for small clusters, but the resonance becomes less evident for even larger clusters, *e.g.*, more than 10 AuNPs. This effect is expected to be related to inter-particle coupling. It also indicates that the experimentally observed extinction is likely mainly attributed to small clusters and AuNP monomers.

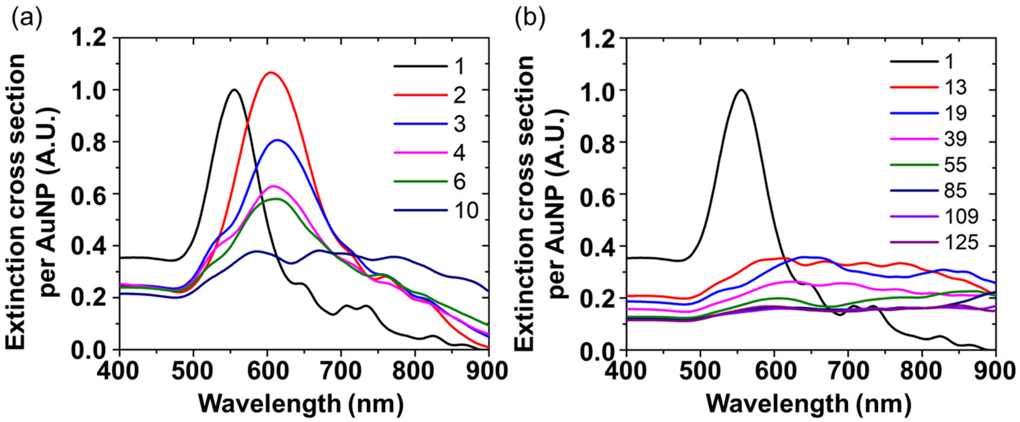


**Figure S2. FDTD-simulated extinction of AuNP clusters**. Extinction cross section per AuNP for 80 nm AuNP clusters with AuNP number of 1-10 in (**a**) and 13-125 in (**b**). Extinction cross section is normalized to AuNP monomer.

#### 3. Impact of nanoparticle size: sGP sensing by incubation

Here, the AuNP size effect was studied with NP diameters of 40, 60, 80, and 100 nm in sensing of Ebola sGP proteins from 1 pM to 1 μM in 1× PBS buffer. The sGP signals were collected using a UV-visible spectrometer coupled to an upright microscope (Figure S3a). We custom designed a polydimethylsiloxane (PDMS) well plate, consisting of 2 mm diameter and 3 mm thick punched holes, that is bonded to a 0.5 mm thick diced fused silica (Figure S3b).

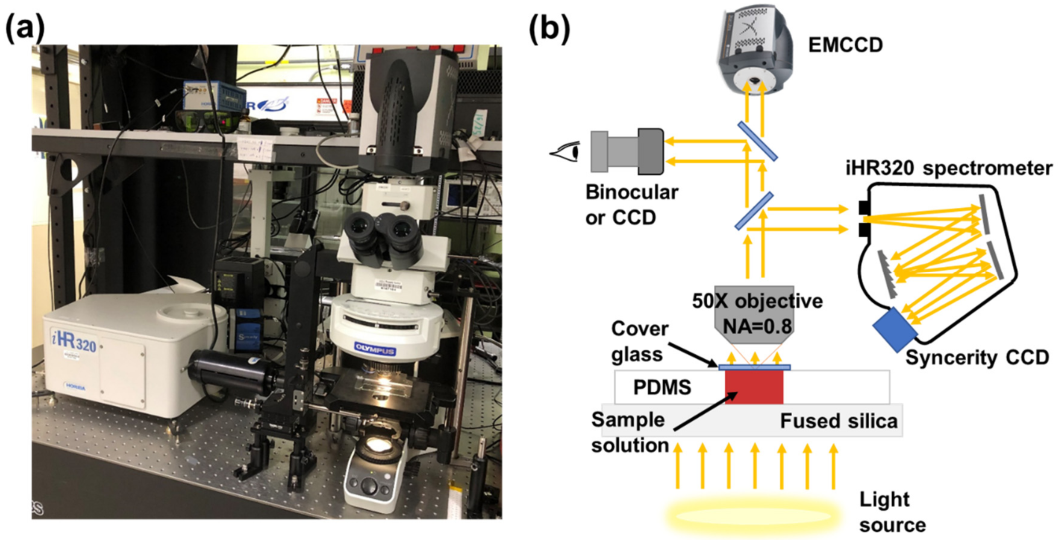


**Figure S3. Experimental setup for spectrometer measurement.** (**a**) Visual image of our lab-based UV-visible spectrometer system consisting of a Horiba iHR320 spectrometer and Olympus BX53 microscope. (**b**) Schematic of the characterization setup for AuNP colloid-based assay using PDMS well plate as a sample cuvette.

Additionally, the AuNP concentrations were adjusted to have roughly identical optical density levels at their peak plasmonic resonance wavelengths (533, 544, 559, and 578 nm for 40, 60, 80, and 100 nm diameter), at an AuNP concentration $[NP]$ of 0.275, 0.086, 0.036, and 0.019 nM, respectively. The extinction coefficient of NPs is theoretically proportional to their total mass (or volume) as $\sigma_{ext} \propto [NP]d^3$ [1], therefore, $[NP]$ drops with the particle diameter given we intentionally standardize the total extinction of all the NPs.

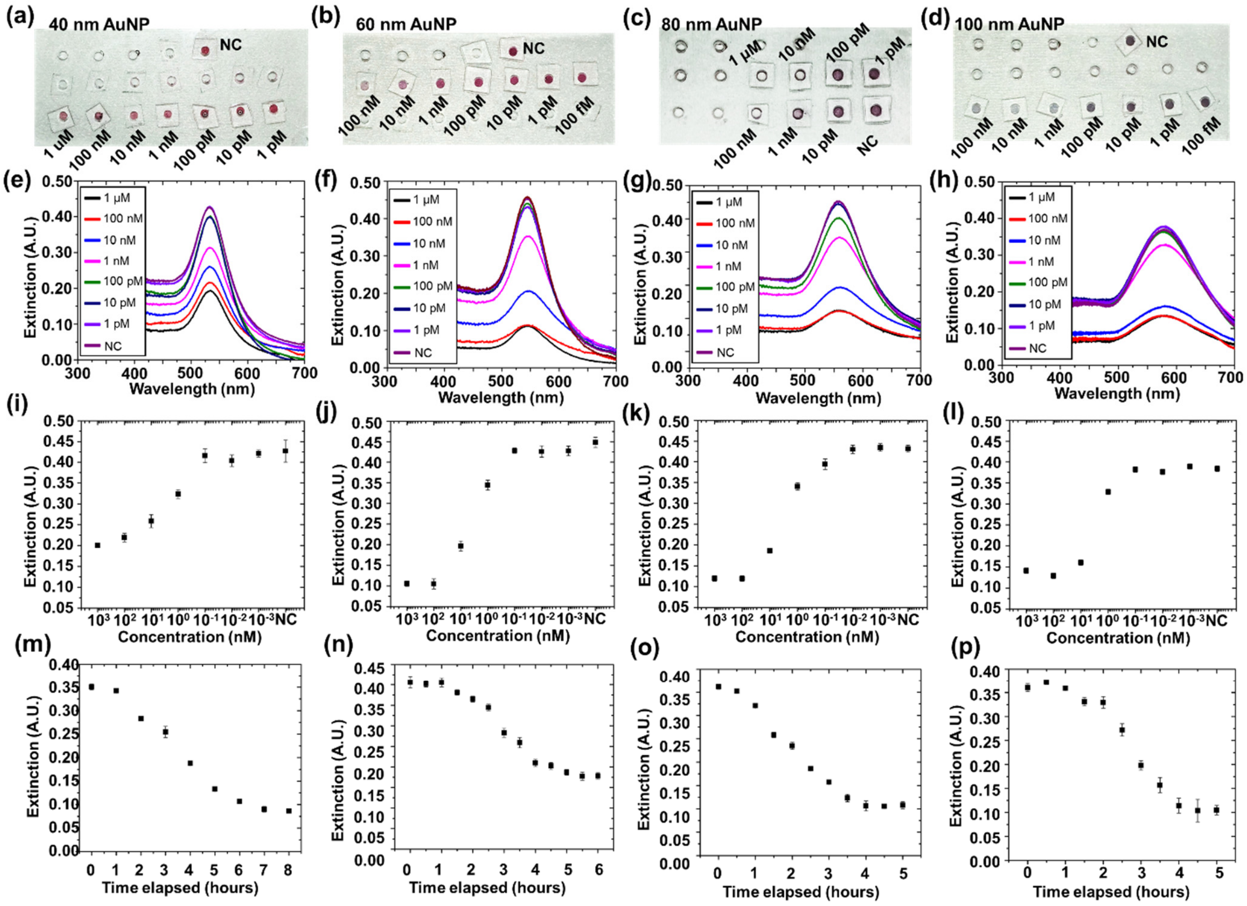


**Figure S4. Study of the impact of nanoparticle size on sGP sensing performance in 1 × PBS buffer.** (**a**)-(**d**) Visual image of 40, 60, 80, and 100 nm AuNP assay upper-level liquid samples loaded on a PDMS well plate, 8, 4, 3, and 3 hours after AuNP assay mixing with 1 pM to 1 μM sGP in 1 × PBS, respectively. Concentrations were assigned in logarithmic scale. Negative control (no sGP) is labeled as NC in all the panels. (**e**)-(**h**) Extinction spectra of samples shown in (**a**)-(**d**) panel, measured by a lab-based UV-visible spectrometer system. The extinction spectra were normalized to transmission through well plate with the same buffer but without any antigens or NPs. (**i**)-(**l**) Extinction peak values plotted for AuNP assay using different NP sizes in detecting sGP with concentration from 1 pM to 1μM. Extinction values are derived from the peak extinction values in (**e**)**-**(**h**). (**m**)-(**p**) Time-dependent response for AuNP assay of different sizes. The sGP concentration was 10 nM.

Here the UV-visible extinction (Figure S4 e-h) can be mathematically defined as $E = \log_{10}\left(\frac{I_0}{I}\right) = \varepsilon c l$, where $E$ is the measured AuNP extinction, $I_0$ and $I$ are the light intensity collected at the reference and sGP sample cuvettes, $\varepsilon$ is the AuNP extinction coefficient, $c$ is AuNP concentration, and $l$ is optical path (the solution depth, ~3 mm here).

We further extracted the extinction peak intensity for each assay sample and plotted the standard curves of extinction versus concentration for each AuNP size (Figure S4 i-l). The sGP signal at low concentration (<10 pM) was indistinguishable from the NC signal (with an extinction $E_{NC}$ within the range of 0.4-0.45).

In addition to the detection limits, the assay detection time was studied at 10 nM sGP concentration in 1× PBS (Figure S4 m-p). The extinction generally started to drop after 0.5-1.5 hour for all sizes, indicating a stage to initiate aggregate formation and precipitation. Extended incubation led to a nearly linear extinction drop, at a rate of 0.049, 0.071, 0.080, and 0.091 $hr^{-1}$ for 40, 60, 80, and 100 nm NPs, eventually reaching a stable value after 7, 5.5, 4, and 4.5 hours, respectively.

### 4. Characterization of sGP sensing monobinders

#### 4.1 Preparation of mono-binder surface functioned AuNP colloid.

The gold nanoparticles (0.13 nM, 80 µL) that were already surface-functioned with streptavidin were first mixed with an excessive amount of biotinylated sGP49 nanobody (1.2 µM, 25 µL). The mixture was then incubated for 2 hours to ensure complete streptavidin-biotin conjugation. Next, the mixture was purified by centrifuge (accuSpin Micro 17, Thermo Fisher) at 10,000 rpm for 10 mins and repeated twice to remove unbounded biotinylated sGP49 nanobody. The purified AuNP colloid was measured by Nanodrop 2000 (Thermo Fisher) to determine the final concentration. The concentration of AuNP in colloid was subsequently adjusted to 0.048 nM and was aliquoted into 12 uL in a 500 uL Eppendorf tube. sGP stock solution (6 µM, in 1×PBS) underwent a 10-fold serial dilution and a 4 uL sGP solution of each concentration (4 pM to 4 uM) was mixed with 12 uL AuNP assay colloid and briefly vortexed (mini vortexer, Thermo Fisher) at 800 rpm for 5 seconds. The buffer used in assay preparation and sGP dilution was prepared by diluting 10×PBS buffer and mixing with glycerol and BSA to reach a final concentration of 1×PBS, 20% v/v glycerol and 1 wt% BSA.

#### 4.2 TEM inspection.

The assay colloids at the bottom of Eppendorf tube were collected and imaged by cryogenic transmission electron microscope (CryoTEM). Cryogenic sample preparation is known to prevent water crystallization and hence preserve protein structures and protein-protein interaction. Clearly, aggregates of AuNPs formed with 1 µM sGP present in the precipitates with average size of 2.3 by 1.4 µm. The cluster size averaged 1.8 by 1.4 µm with 1 nM sGP.

For the NC sample, only 80 nm AuNP monomers but no clusters were observed in the precipitates (Figure S5). The aggregate sizes had a large distribution, probably due to a random aggregation process and sample preparation, and thus was not ideally correlated with the sGP concentration.

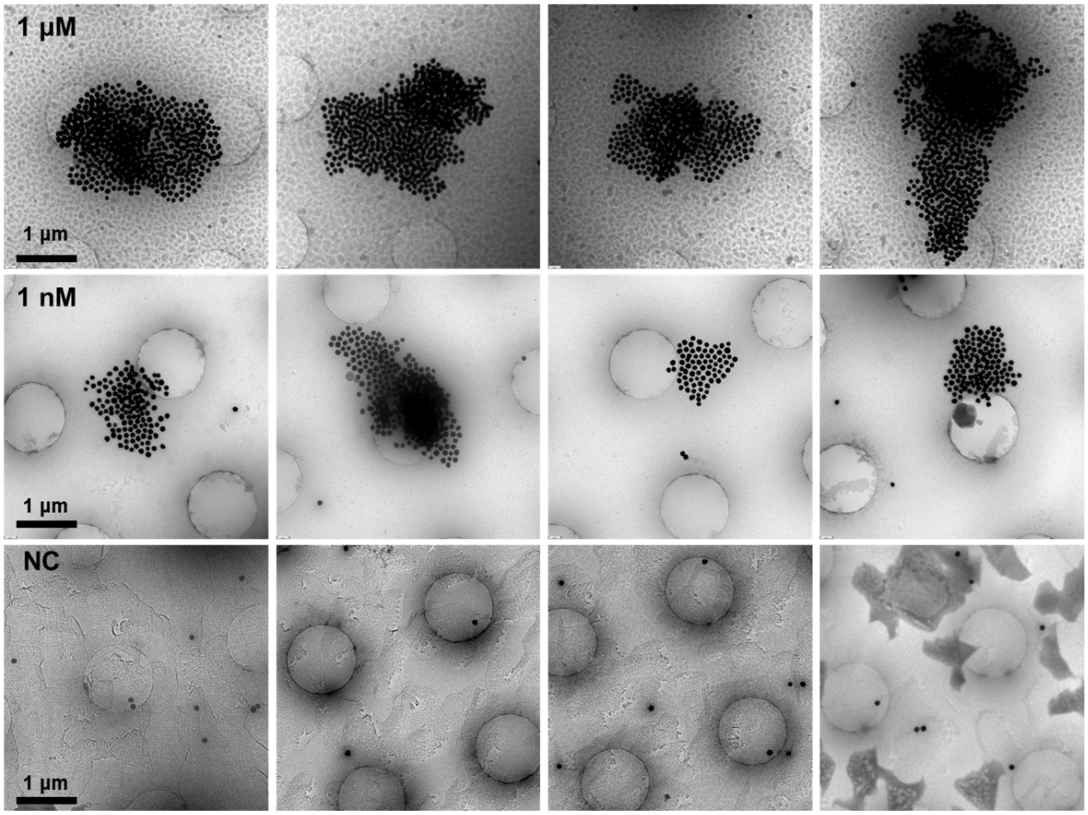


**Figure S5. Representative cryoTEM images of 80 nm AuNP assay in detecting 1 nM, 1 μM sGP, and NC samples.**

**4.3 Drop-casting on glass slides for optical inspection**

The AuNP assay upper-level liquid (1 μL, sGP mono-binder, 80 nm AuNP, $n_{AuNP}$ = 0.036 nM) with sGP from 1 pM to 1 μM in PBS were drop-casted on a 1 mm thick glass slide for colorimetric and spectrometric inspections (Figure S6). It can be observed that the dried sample spots displayed light red color, and their transparency increased from around 10 nM (spot 3) and became readily differentiable to the naked eye at 1 μM (spot 1) compared to the reference NC sample (spot 8). We then measured the extinction spectra of each drop-cast spots and extracted the extinction peak intensity at LSPR resonance (Figure S6c). The extinction spectra featured AuNP LSPR peaks, similar to those upper-level liquid measurements in the PDMS well plate, but the peak intensity was about one order of magnitude smaller attributed to a significantly shorter optical path (estimated ~300 μm) compared to the PDMS well plate (~3 mm).

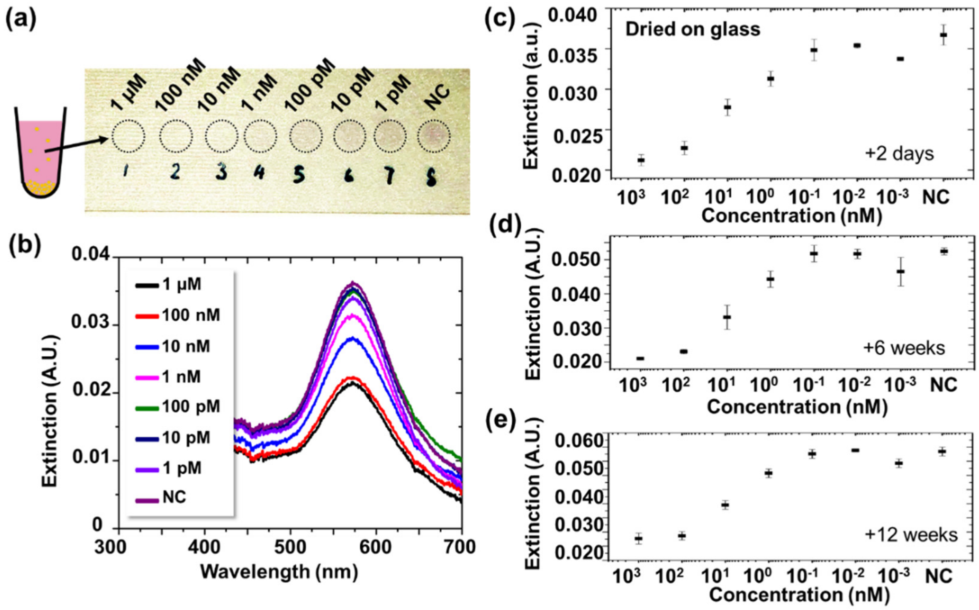


**Figure S6. sGP protein sensing by drop-cast on a glass slide.** (**a**) Visual image of AuNP assay upper-level liquid dried on a 1 mm thick glass slide. (**b**) Extinction spectra of drop-cast samples shown in (**a**), measured by UV-visible spectrometer. The extinction spectra were normalized to transmission through bare glass slides without any biological samples. (**c**) Extinction peak values (at 573 nm) extracted from Figure (b), 2 days after drop casting. (**d-e**) Extinction peak values measured 2 days after drop casting. Here, 80 nm AuNP were functionalized with sGP49 on 80 nm AuNPs. Negative control (no sGP) is labeled as NC for comparison. In the test, sGP concentration ranges from 1 pM to 1 μM.

These samples were stored at room temperature (25 ˚C) over 12 weeks and re-inspected, and the optical signals were found in general to be consistent over such an extended period with only slight change (Figure S6). The slight increase in extinction, observed especially at the lower sGP concentration, was possibly due to shrinking of the drop cast spot from dehydration as the sample was exposed to dry air. Nevertheless, this also showed the feasibility of quantitative detection of sGP down to 350 pM LoD with a broad dynamic range in detection (100 pM to 1 μM) using a simple and small solid-state sample carrier.

In comparison, we also performed drop-casting of 60 nm AuNPs on glass slides (Figure S7), which also shows performance similar to the detection in the PDMS well plate (Figure 2).

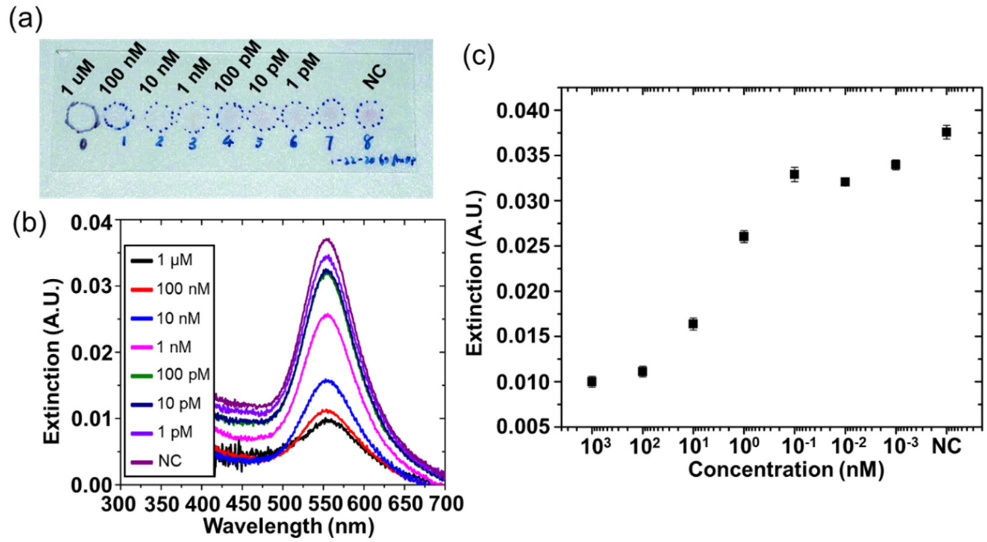


**Figure S7. UV-visible spectrometer characterization of 60 nm AuNP drop cast assay.** (**a**) Visible image of drop cast samples on a 1 mm thick glass slide. (**b**) Extinction spectra measured for samples shown in (**a**). The extinction spectra were normalized to transmission through bare glass slides without any biological samples. (**c**) Extinction maximum at AuNP resonance wavelength, extracted from extinction spectra shown in (**b**).

#### 4.4 Drop-casting on gold film for SEM and dark-field inspection

Scanning electron microscopy (SEM) was employed to investigate the AuNP aggregation in the upper-level liquid. Here 1 µL of 80 nm AuNP assay colloids ($n_{AuNP}$ = 0.036 nM, 1×PBS) with targeted sGP (1 pM to 1 µM) were drop-cast onto an oxygen plasma treated gold surface and subsequently dried in air (Figure S8). Gold surface was selected due to its high electron conductivity that dramatically improves the contrast and resolution in imaging. Only AuNP monomers but no large aggregates were observed from the SEM images, confirming that the majority, if not all, of the aggregates should precipitate at the bottom of the tubes. Further, the 80 nm AuNPs were recognized and counted through image analysis, and their density was statistically determined from ten SEM images (total area 10 × 8.446 × 5.913 $\mu m^2$) at each sGP concentration (Figure S8 c). Clearly, the AuNP density decreased at a higher sGP concentration, i.e. from 1.97 $\mu m^{-2}$ at about 10 pM to 0.26 $\mu m^{-2}$ at 100 nM and finally saturated to 0.24 $\mu m^{-2}$ at about 1 µM or above, which was in accordance with extinction spectrometric measurements of both upper-level liquid samples and glass slide drop-cast samples. The limit of detection derived from SEM characterization was estimated to be about 150 pM, comparable but slightly higher than the upper-level liquid extinction characterization for the same 80 nm AuNPs in PBS, possibly due to increased variance in nanoscale level characterization and limited sampling data.

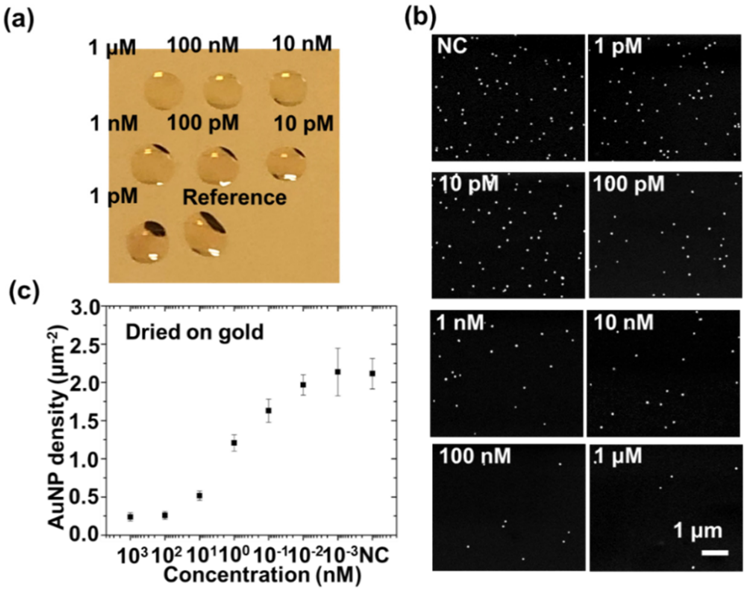


**Figure S8. sGP protein sensing by drop-cast on gold film.** (**a**) Visual image of AuNP assay upper-level liquid dried on a 100 nm gold film thermally evaporated on a silicon substrate. (**b**) Representative SEM images of drop-cast samples shown in (**a**). (**c**) AuNP surface density measured from SEM images (10 images used for each data point, including those shown in (**b**)). In the test, sGP concentration ranged from 1 pM to 1 μM. Here, 80 nm AuNP were functionalized with sGP49 on 80 nm AuNPs. Negative control (no sGP) is labeled as NC for comparison.

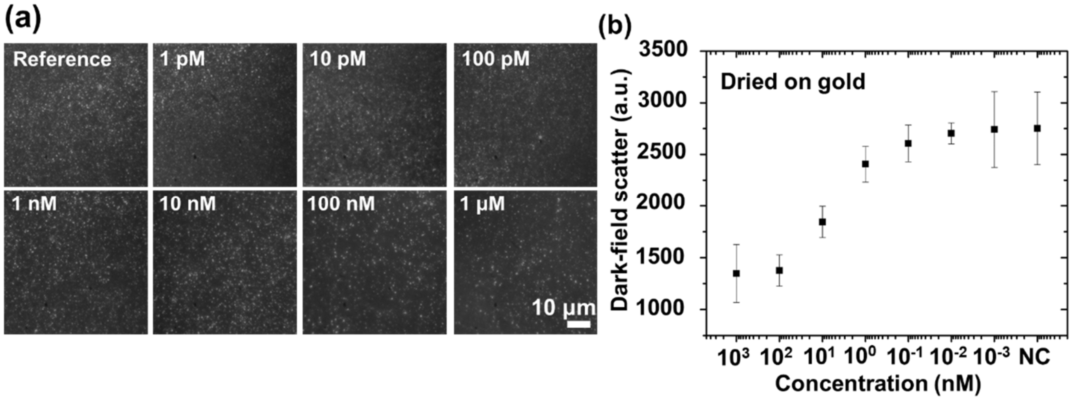


**Figure S9. Dark field scattering image of 80 nm AuNP assay samples drop cast on Au surface.** (a) Representative dark-field images at different concentrations. (b) The counted dark-field signal spots at different concentrations.

Further, the drop-cast samples on gold surface were also analyzed by dark field scattering imaging. Here LSPR mediated scattering of incident light from AuNPs on gold surface can directly indicate the density of AuNP, based on the density of bright spots on dark field imaging (Figure S9). The dark field scattering images were undergoing image process to improve the contrast for spots, which were then counted using MATLAB code and averaged over 10 images (captured area 62.5 μm × 62.5 μm in each image). It was observed that the density of bright spots dropped as sGP concentration increased, consistent with SEM observation. The dark field imaging method was estimated to be capable of detecting sGP with a sensitivity about 1 nM.

#### 4.5 Drop-casting on gold film in detecting sGP in 1 × PBS with 100 nm AuNPs

To investigate the feasibility of different AuNP sizes on sensing, we prepared 100 nm AuNP assay samples, drop-cast them on 1mm glass slides and gold films, and characterized by UV-visible spectrometer, SEM, and dark field scattering imaging, as shown in Figure S10. The 100 nm AuNP concentration in assay colloid was 0.019 nM. The characterizations followed protocols described in main context method: UV-visible spectra, SEM imaging, and dark field scattering imaging characterizations. The measurement results in general showed comparable sensitivity (sub 1 nM) to that of 80 nm AuNPs and consistent with detection in PDMS well plate.

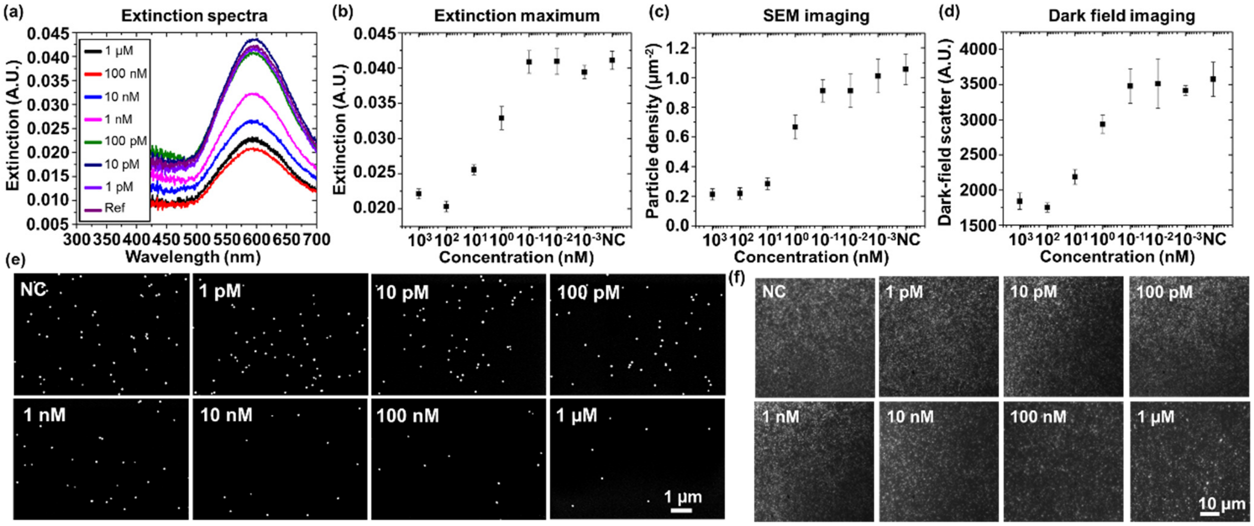


**Figure S10 Characterizations of drop-casted 100 nm AuNPs.** (**a**) Extinction spectra measured for 100 nm AuNP drop cast samples. (**b**) Extinction maximum at AuNP resonance wavelength, extracted from extinction spectra shown in (**a**). (c) Particle density average calculated for ten SEM images taken of 100 nm AuNP drop cast samples. (**d**) Scattered particle counts statistically derived for ten dark field scattering images taken for 100 nm AuNP drop cast samples. (e) SEM image of 100 nm AuNP assay samples drop cast on Au surface. (**f**) Dark field imaging.

### 5. Back-of-the-envelope estimation to understand the detection limits

Our experimentally determined upper and lower limits in detection are thought to be correlated to the multivalent binding nature of the detection process. Suppliers informed that up to 120, 270, 460, and 730 streptavidin are bound on 40, 60, 80 and 100 nm diameter NPs, although the effective numbers are expected to be smaller. This large number of binding sites on the AuNPs makes them a strong multivalent binding sensor that is highly favorable to AuNP aggregation, even at low antigen concentration.

With very small amounts of antigens, the ultimate lower limit of detection occurs when NP precipitation decreases the AuNP monomers in the suspension significantly enough to produce a signal that is distinguishable from noise or fluctuation. This value is expected to be dependent on both the dynamic antigen-antibody binding process and the experimental setup. For example, assuming a very high affinity in sGP binding (ignoring dissociation) and four sGP bound to each aggregating AuNP (about 3-5 at 1 nM from TEM image from Figure S5) and setting 3% optical extinction change as the detection threshold to overcome signal variation, we could roughly estimate the LoD as 275×(3%/45%)×4=70 pM or 36×(3%/45%)×4=10 pM for 40 and 80 nm AuNPs, which are comparable to our experimental analysis. Similarly, the upper limit of detection could be estimated when the AuNPs are completely saturated with the analyte, *i.e.* 120×0.275=33 nM and 460×0.036=16 nM for 40 and 80 nm AuNPs, also comparable to but smaller than experimental values. The above back-of-the-envelope analysis is helpful to provide intuitive understanding of the measured dynamic range and detection limits, but it is also quite limited because it ignores the dynamic association and dissociation processes which are thought to be dependent on both the NP size and antigen-nanobody binding characteristics.

### 6. Modeling to understand sensing physics

#### 6.1 Rate limiting reaction steps and diffusion

Using 80 nm AuNP as an example, we first attempted to identify the key rate determining step in the sensing mechanism. Each of the 80 nm AuNPs has ~460 nanobodies on their surface and behaves as a multivalent sGP-binding pseudo-particle that diffuses and conjugates to each other via sGP-mediated bridging. This triggers formation of AuNP dimers oligomers, and eventually large clusters, which precipitate at the bottom of microcentrifuge tube as gravity gradually overtakes fluidic drag force. Therefore, the reaction determining steps during the sensing process include the antigen diffusion, AuNP diffusion, antigen-AuNP binding, AuNP clustering, and AuNP precipitation.

The diffusivities of AuNPs and antigens can be estimated from the Stokes−Einstein equation $D = kT/(3\pi\eta d)$, where $kT$ is the thermal energy at room temperature (~$4.1\times10^{-21}$ Joule at 300K), $\eta$ is the solution viscosity (~$1.7\times10^{-3}$ N·sec/m$^2$ assuming 20% glycerol in water to estimate the buffer effect), and $d$ is the particle diameter[2]. The diffusivity is estimated $D_a$~$5.12\times10^{-11}$ m$^2$/s for a 5 nm protein and $D_{NP}$ ~$3.2\times10^{-12}$ m$^2$/s for an 80 nm AuNP. We can further estimate the diffusion length $L_a$, *i.e.* the distance for analyte to collide with AuNPs, as the smaller of the inter-protein separation $L_p$ and inter-AuNP separation $L_{NP}$. $L_{NP}$ was calculated in the range of 2 to 5 μm for 40 to 100 nm AuNPs used in our experiment following $L_{NP} = \sqrt[3]{\frac{6V}{\pi}} = \sqrt[3]{\frac{6}{\pi}\frac{1}{c_{NP}N_A}}$, where $V$ is volume estimated for each NP (assuming a sphere), $c$

is the NP molar concentration, and $N_A$ is the Avogadro number. Clearly, $L_{NP}$ is determined by the NP concentration and thus is a constant once the assay is designed. Similarly, $L_p$ depends on analyte concentration and can be calculated as ~2 μm at a low sGP concentration (<100 pM) but <100 nm at a higher concentration (>1 μM). Therefore, $L_a$ is mainly determined by the protein concentration, and the diffusion time $t_a \sim L_a^2/D_a$ is found to be only 0.1 to 0.2 sec, much shorter than the experimentally determined incubation assay time (3 to 7 hours, Figure S5).

It is important to compare here with conventional surface-incubation based assays, such as ELISA and SPR, where according to the non-slip boundary condition the surface velocity is close to zero. Differently, in our case the AuNP continues to diffuse, and its thermal velocity can be estimated by $v_{NP} = \sqrt{KT/m_{NP}}$. Because of its small mass ($m_{NP} = 5.18 \times 10^{-15} g$ for an 80 nm AuNP), there is a significantly large thermal velocity for the AuNPs $v_{NP}$~0.028 m/sec. Given this velocity and the small $L_a$, we can understand that in fact the AuNPs should be constantly colliding with antigens and other nanoparticles, promoting effective mixing and antigen binding. Therefore, the diffusion process will not be a rate limiting step here, although they could significantly limit the assay time of ELISA and SPR. In another word, the antigen binding process behaves totally differently from that on an infinitely large surface in ELISA and SPR, and the measured $k_{on}$, $k_{off}$ from ELISA is limited to the surface bound molecular interactions and cannot fully predict what happens at the nanometer scale in our case. Such phenomena have been observed that the binding kinetics in solution could be significantly different from that on surface [3], which is attributed to mass transport and other factors[4]. On the other hand, given the high binding affinity of analyte-ligand complex in this proposed work, we can reasonably hypothesize that the association process of this complex is fast (*e.g.,* G protein binds to GPCR receptors within ~0.3 sec[5]), thus also unlikely the limiting step.

**6.2 Simplified mathematical modeling of AuNP aggregation**

Here we adapted Smoluchowski's coagulation equation and modified the equation to describe our reversible AuNP aggregation process, using sGP sensing as an example. The modified equation is:

$$\frac{dn_i}{dt} = \frac{1}{2}\sum_{j=1}^{i-1} k_{i-j,j} n_{i-j} n_j - \sum_{j=1}^{\infty} k_{i,j} n_i n_j + k_{off} n_{i+1} - k_{off} n_i \quad (1)$$

Where $n_i(t)$ is the concentration of aggregates consisting of $i$ AuNPs, $k_{i,j}$ is the coagulation kernel for the aggregation of clusters consisting of $i$ AuNPs and $j$ AuNPs, $k_{off}$ is the dissociation constant of sGP49-sGP conjugation ($k_{off}$=1.88×10$^{-4}$ s$^{-1}$) derived from ELISA measurement. According to Brownian diffusion theory, the coagulation kernel $k_{i,j}$ is described as [6, 7]:

$$k_{i,j} = \frac{2}{3} P \frac{k_B T}{\eta} (ij)^{\gamma} \left( m_i^{\frac{1}{d_f}} + m_j^{\frac{1}{d_f}} \right) \left( m_i^{-\frac{1}{d_f}} + m_j^{-\frac{1}{d_f}} \right) \quad (2)$$

Where $P$ is the probability of aggregation per collision, $k_B$ is Boltzmann constant, $T$ is temperature, $\eta$ is dynamic viscosity of colloid buffer (~$1.7\times10^{-3}$ N·s/m$^2$ for 20% glycerol in water), $i$ and $j$ are numbers of AuNP in each cluster, $m_i$ and $m_j$ are mass of each clusters, $d_f$ is the fractal dimension (~2.1 for a typical densely aggregated cluster). Here $P$ could be estimated as 0.0165 if using the ELISA-determined kinetic parameter $k_{on}$=$4.07\times10^4$ M$^{-1}$s$^{-1}$. This value is found a serious underestimate because the mass transport and antigen binding process in in-solution Nano2RED assay are significantly more effective than that for ELISA, as discussed in section 6.1. Indeed, we found using such a small $P$ value could not accurately predict the assay time observed in Nano2RED. Instead we used $P$=1, meaning every collision of sGP and sGP49-functionalized AuNPs will result in antigen binding. Such an assumption led to good agreement between theory and experimental observations. In fact, such a high binding efficiency is reasonable given the multivalence nature of AuNP sensors. The multivalence effectively creates a much higher "functional affinity" compared to the intrinsic affinity by monovalent binding[8, 9], and could yield virtually irreversible binding process.

In equation 1, the first two terms in the right side directly come from Smoluchowski's equation that describe the AuNP aggregation process. The other two terms on the right side are added terms to describe the reversible dissociation of AuNP aggregates. For simplification, we considered only the dissociation of a cluster with N AuNPs to form a cluster of N-1 AuNPs and a monomer released back to colloid (*i.e.* N → N-1,1). Although the breakdown of clusters to clusters with other arbitrary numbers of AuNPs is possible (N → N-i, i), such breakdown requires multivalent sGP-sGP49 dissociation, hence the effective dissociation rate is likely to be much smaller. Moreover, a further simplification of the model by considering only the low-order oligomers and monomers interactions is justified by the fact that the concentration of $n_i$ with higher $i$ numbers (higher order) is small due to precipitation. Therefore, we could calculate the AuNP monomer concentration based on the simplified equation set and thus estimate the extinction signals by considering only the low-order oligomer-monomer interactions. This assumption is especially valid at the beginning when the assay is mixed with sGP protein, where the concentration of higher order oligomers and large clusters is near 0. Figure 5**a** shows the schematic of low-order oligomer-monomer interaction and the evolution of large clusters formation. Our simplified model incorporates such evolution, and the equation sets are shown below:

$$\frac{dn_1}{dt} = -2k_{11}{n_1}^2 - \sum_{i=2}^{\infty} k_{1i}n_i n_1 + 2k_{off}n_2 + \sum_{i=3}^{\infty} k_{off}n_i \qquad (3)$$

$$\frac{dn_2}{dt} = 2k_{11}{n_1}^2 - k_{12}n_1 n_2 + k_{off}n_3 - 2k_{off}n_2 \qquad (4)$$

$$\frac{dn_3}{dt} = k_{12}n_1 n_2 - k_{13}n_1 n_3 + k_{off}n_4 - k_{off}n_3 \qquad (5)$$

…

$$\frac{dn_i}{dt} = k_{1i-1}n_1n_{i-1} - k_{1i}n_1n_i + k_{off}n_{i+1} - k_{off}n_i \qquad (6)$$

By solving the equations above, we obtained the time-dependent monomer concentration, assuming 0.036 nM AuNP colloid in detecting 10 nM sGP, and further converted the concentration to optical extinction, as shown by the black curve in Figure 5**b**. Intuitively, the solver of this equation set showed that the monomer concentration versus time is quasi-exponential. The aggregation time constant $\tau_{agg}$, defined as time required for concentration of AuNP monomer to drop to $c_{equillibrium} + \frac{1}{e}(c_0 - c_{equillibrium})$, is 0.87 hour.

Further, we calculated the monomer concentration versus time for 1.8 nM 80 nm AuNP assay in detecting 10 nM sGP. The time-dependent extinction is shown by the red curve in Figure 5**b**. In this case, $\tau_{agg}$ is significantly shortened to 0.024 hour, or ~36 times smaller compared to $\tau_{agg}$ using 0.036 nM AuNPs.

#### 6.3 Sedimentation time

As AuNPs cluster, the gravitational force overcomes fluidic drag, and large clusters precipitate to form sedimentation and continuously deplete AuNPs and sGP proteins in the colloid until reaching equilibrium. The sedimentation time of this progression can be estimated from the solution of the Mason-Weaver equation by $\tau_{sed} = z/(s \cdot g)$ where $z$ is the precipitation path (the height of colloid liquid), $g$ is the gravitation constant, $s$ is the sedimentation coefficient $s = \frac{d^2}{18\eta}(\rho_M - \rho_w)$ ($d$ is the aggregate diameter, $\eta$ is the dynamic viscosity of the colloid buffer, $\rho_M$ and $\rho_w$ are the density of aggregate and colloid buffer, respectively) [10]. Clearly, the large density contrast between gold (19.3g/cm$^3$) and buffer (~1g/cm$^3$) is also beneficial to improve the sedimentation coefficient $s$. Given $z$ ~3.5 mm for 16 μL liquid in a microcentrifuge tube, we calculated that $\tau_{sed}$ decreases from 26.0 hours for an 80 nm AuNP monomer to 1.0 and 0.3 hours for 400 nm and 800 nm diameter clusters, respectively.

### 7. Rapid Detection

#### 7.1 Impact of incubation time

The optical images of the microcentrifuge tubes (Figure S11**a**) indicated the color contrast was high enough to be immediately resolved by the naked eye after vortex mixing. We have analyzed the peak extinction of the assay upper-level liquid at different incubation times, and the extinction was found to be 0.145 right after vortex, completely distinguishable from the NC sample (0.536), and it gradually decreased to 0.110 as incubation time was extended to 20 min. The analysis show that further incubation could be used as an option to moderately improve the signal contrast, but could possibly be skipped when testing speed is extremely important.

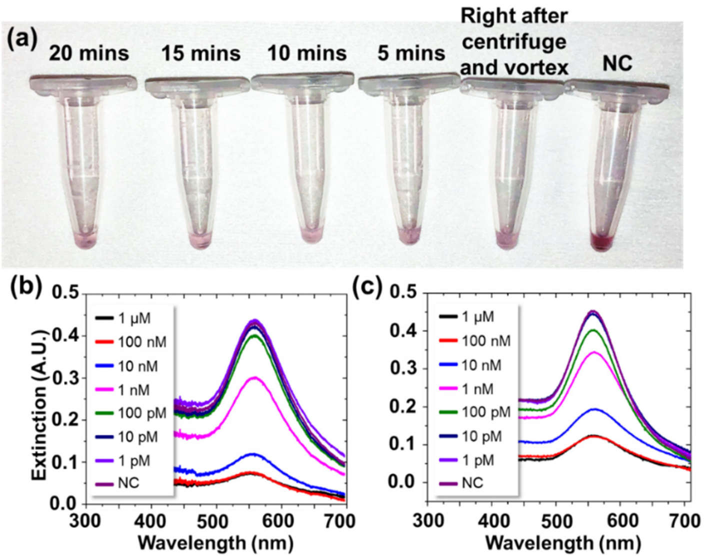


**Figure S11. Centrifuge enhanced 80 nm AuNP assay detecting 10 nM sGP in 1×PBS with different incubation times.** (**a**) Visual images of microcentrifuge tubes. (**b**) Extinction spectra of assay upper-level liquid in PDMS well plate after 20 minutes incubation time using centrifuge-enhanced rapid test. (**c**) Extinction spectra of assay upper-level liquid in PDMS well plate after 3 hours incubation time, without using centrifuge to accelerate readout.

In addition, the rapid sGP detection scheme was found to be reproducible in fetal bovine serum (5% FBS; Figure S12 **a-b**), and it produced similar results in 1×PBS buffer (Figure S12 **c-d**). In both cases, sGP >1 nM can be accurately read out by the naked eye either in tubes or in a PDMS well plate.

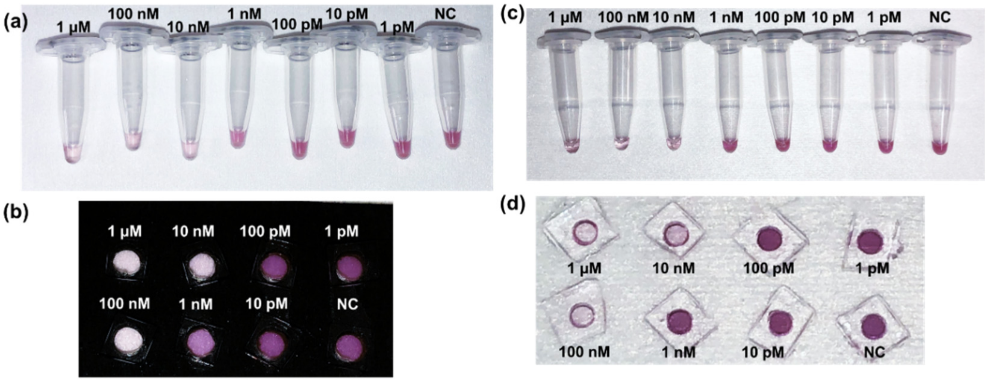


**Figure S12. Visual images for 80 nm AuNP assays detecting 1 pM to 1 μM sGP in 5% FBS (a-b) and 1×PBS (c-d).** Test in 5% FBS: (**a**) optical images of microcentrifuge tubes, with 1 min centrifugation and 15 seconds vortex. Assay upper-level liquid in (**a**) were loaded in 4 mm wells on a 3mm thick PDMS well plate, as shown in (**b**). The region in absence of assay was covered by black flocked paper (Thorlabs, BFP1) that absorb 99% of light from 420 nm to 650 nm. Test in 1×PBS: (**c**) optical images of microcentrifuge tubes. Assay upper-level liquid in (**c**) were loaded in 2 mm holes on a 3 mm thick PDMS well plate, shown in (**d**).

#### 7.2 Portable UV-visible spectrometer as readout

The portable UV-visible spectrometer system (Figure S13) consists of a smartphone sized Ocean Optics UV-visible spectrometer (8.8 × 6.3 × 3.1 $cm^3$), a lamp source module (15.8 × 13.5 × 13.5 $cm^3$), alignment clamps, and an electronic recording device (such as a laptop or a smart phone). Here, 80 nm AuNP colloid (30 μL) is mixed with sGP testing buffer (10 μL 5% FBS), vortexed and centrifuged at 1,200 × g (3,500 rpm) for 1 min. After 20 minutes of incubation, the assay colloids were vortexed for 15 seconds and the upper-level liquid were loaded into 4 mm-diameter wells on a 3 mm thick PDMS plate (Figure S12**b**). The light from the lamp transmitted through the colloid, whereas the rest area of the diced fused silica was covered in black to block stray light transmission. Transmitted light was collected by the spectrometer through a fiber waveguide. The extinction spectra, measured by portable spectrometer (Ocean Optics, Figure S13**b**), were in general highly consistent with that measured by microscope-coupled spectrometer (Horiba iHR320, Figure S13**c**), with a slightly increased signal noise. The extinction peak values at the resonance wavelength (~559 nm) of the two measurements were in high agreement, with a small difference within 15.8%. This could possibly be attributed to different signal collection setup (10× objective with NA of 0.3 in lab-based spectrometer versus waveguide collecting a nearly collimated beam in portable spectrometer). The optical signals measured by portable spectrometer were able to distinguish sGP at 100 pM ($E_{100pM}$=0.509) from the reference ($E_{NC}$=0.542, no sGP), with a dynamic range (10 pM to 1 μM) and limit of detection (~42 pM) comparable to that of the rapid detection of sGP in serum and 1 × PBS (Figure 4 and Figure S12).

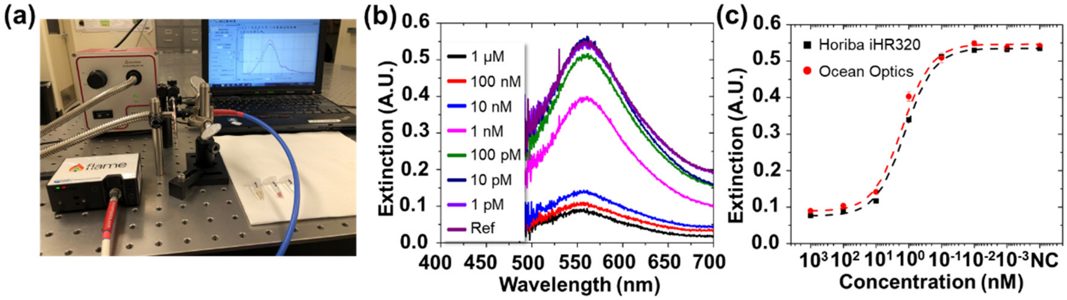


**Figure S13. Detecting sGP in FBS using a portable spectrometer system.** (**a**) Visual image of the measurement system consisting of a laptop (right side of image), lamp source module (left side of image), Ocean Optics Flame spectrometer (right front), and alignment clamps (middle of image). The sample loaded PDMS well plate (visual images shown in Figure S12**b**) is placed in the gap between Ocean Optics spectrometer signal reading waveguide (right) and end of the light source (left). (**b**) Extinction spectra of assay samples in detecting sGP in 5% FBS, measured by Ocean Optics spectrometer. (**c**) Extinction maximum at 80 nm AuNP resonance wavelength $\lambda_p$ =559 nm, derived from spectra in (**b**), shown in red data points. Extinction maxima derived from lab-based spectrometer measured spectra (Figure S3, used 10× objective to collect transmitted light) are shown in black data points for comparison.

### 8. Nano2RED with co-binders and testing in different biological buffers

#### 8.1 The impact of biological buffer concentration

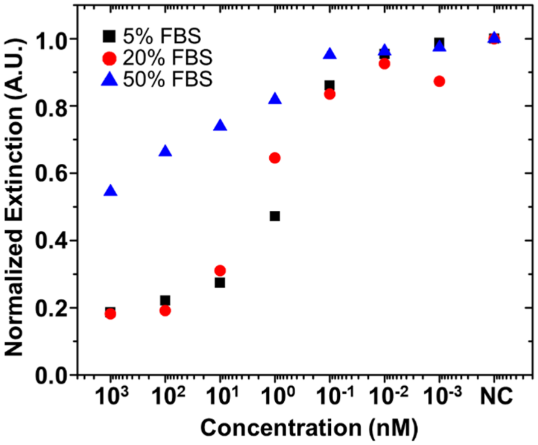


**Figure S14 Normalized extinction for a drop cast 80 nm AuNP assay in detecting sGP in 5, 20, and 50% FBS.** sGP concentrations were spiked from 1 pM to 1 μM.

Using FBS as a buffer and sGP49 functionalized 80 nm AuNPs for sensors, we have investigated the effect of buffer concentration (Figure S14). We found that high concentration FBS (50%) produced smaller optical contrast for readout compared to low concentration, <20% FBS. Further, 5% FBS displayed more consistent results and a slightly larger dynamic range for detection. For consistency in comparison, we chose 5% as the concentration of all biological buffers to be tested in the co-binder experiments.

#### 8.2 sGP detection using co-binder sGP49/sGP7

Two different biotinylated nanobodies, sGP49 and sGP7, were surface functionalized to streptavidin coated AuNP similar to the method described earlier for creating two different sets of functionalized AuNP colloidal solutions. The concentration of functionalized AuNP colloids were re-adjusted to get an optimal extinction level. sGP stock solution underwent serial dilution to create an sGP analyte solution with concentrations of 4 uM to 400 fM in selected detection media. The final composition of PBS detection media was composed of 1×PBS, 20% v/v glycerol and 1 wt% BSA while that of FBS, HPS (Human pooled serum), and WB (Whole blood) detection media had a final concentration of 1×PBS, 20% v/v glycerol and 1 wt% BSA and 20% of either FBS, HPS, or WB which resulted in a final concentration of FBS, HPS, or WB in the detection assay to be 5%. Solutions of sGP49-functionalized AuNPs, sGP7-functionalized AuNP, and sGP were mixed in a 500 uL Eppendorf tube at a ratio of 3:3:2 and thoroughly vortexed. After mixing, the detection assay was centrifuged (accuSpin Micro 17, Thermo Fisher) at 3,500 rpm (1,200×g) for 1 minute. AuNPs were highly concentrated at the bottom of Eppendorf tube. After 20 minutes of incubation, the colloidal assay was vortexed (mini vortexer, Thermo Fisher) at 800 rpm for 15 seconds to thoroughly remix free AuNP monomers into the colloid. Following this, spectrometric and electronic characterizations were done in a way similar to that described previously.

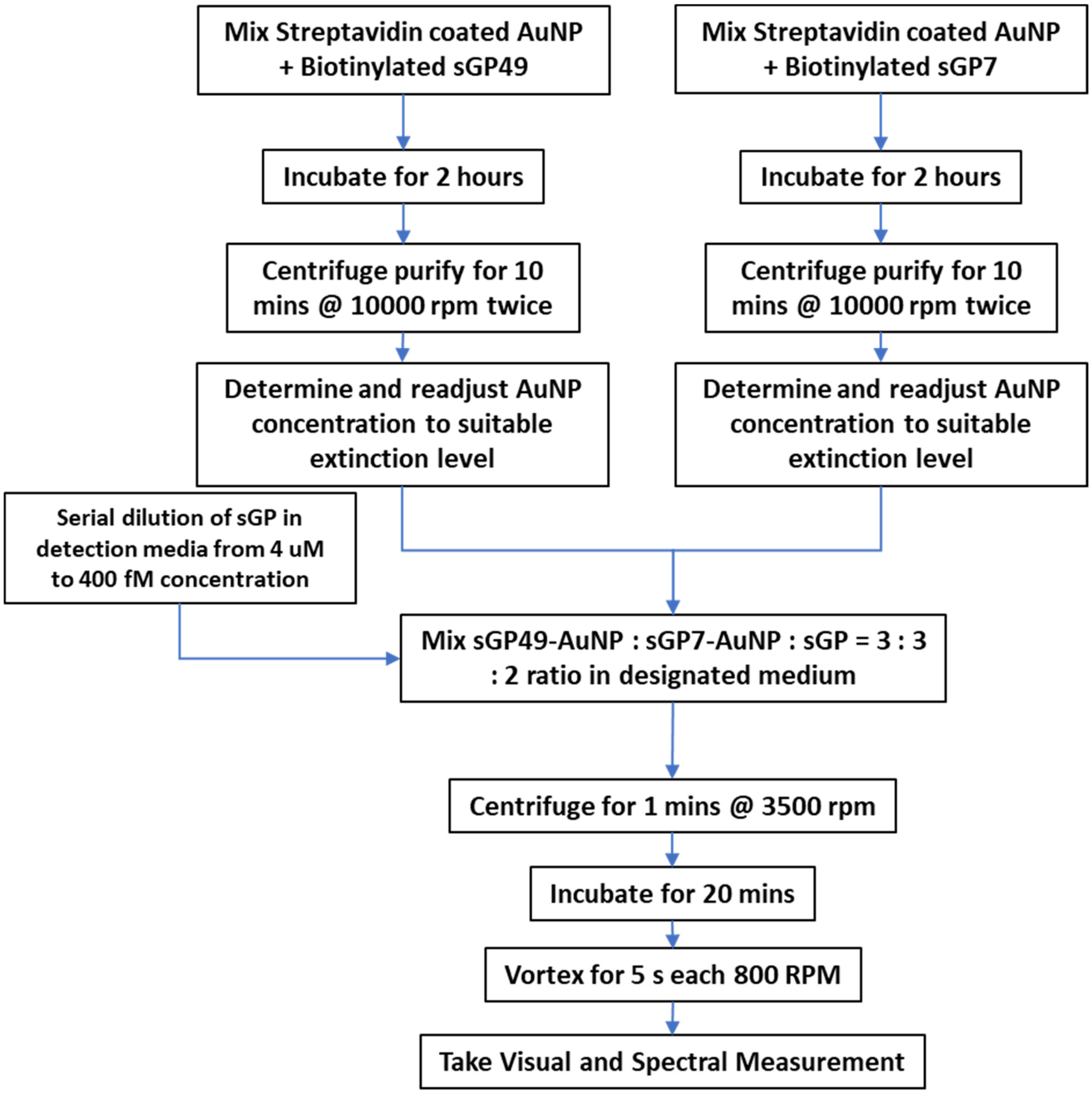


**Figure S15. Workflow of EBOLA antigen sensing using co-binders.**

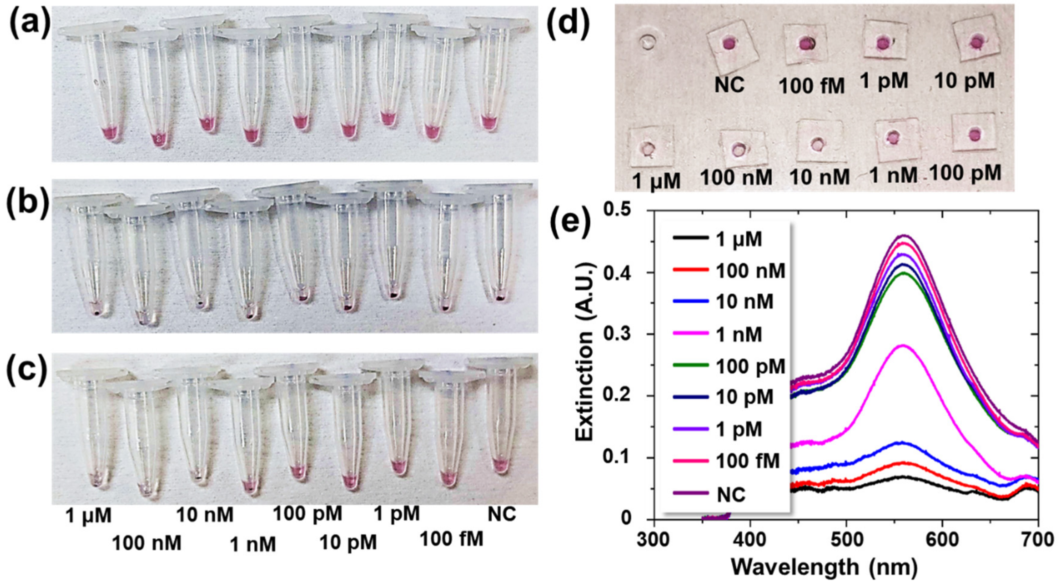


**Figure S16. EBOLA co-binder test in 1X PBS.** (**a-c**) Optical images of microcentrifuge tubes (**a**) after mixing, (**b**) right after centrifugation, and (**c**) after incubation (20 min) and vortex-mixing. (**d**) The upper-level liquid from (**c**) loaded into a PDMS well plate. (**e**) The spectrometric readout from the well plate. The electronic signals shown in Figure 5 were readout from the microcentrifuge tubes after incubation and vortex-mixing.

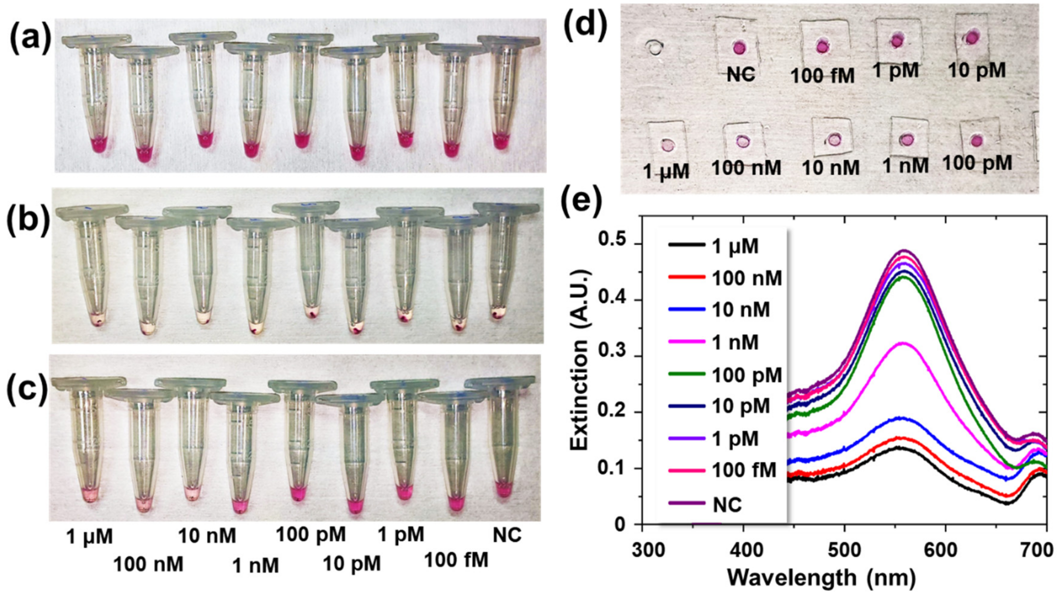


**Figure 17 EBOLA co-binder test in 5% FBS.** (**a-c**) Optical images of microcentrifuge tubes (**a**) after mixing, (**b**) right after centrifugation, and (**c**) after incubation (20 min) and vortex-mixing. (**d**) The upper-level liquid from (**c**) loaded into a PDMS well plate. (**e**) The spectrometric readout from the well plate. The electronic signals shown in Figure 5 were readout from the microcentrifuge tubes after incubation and vortex-mixing.

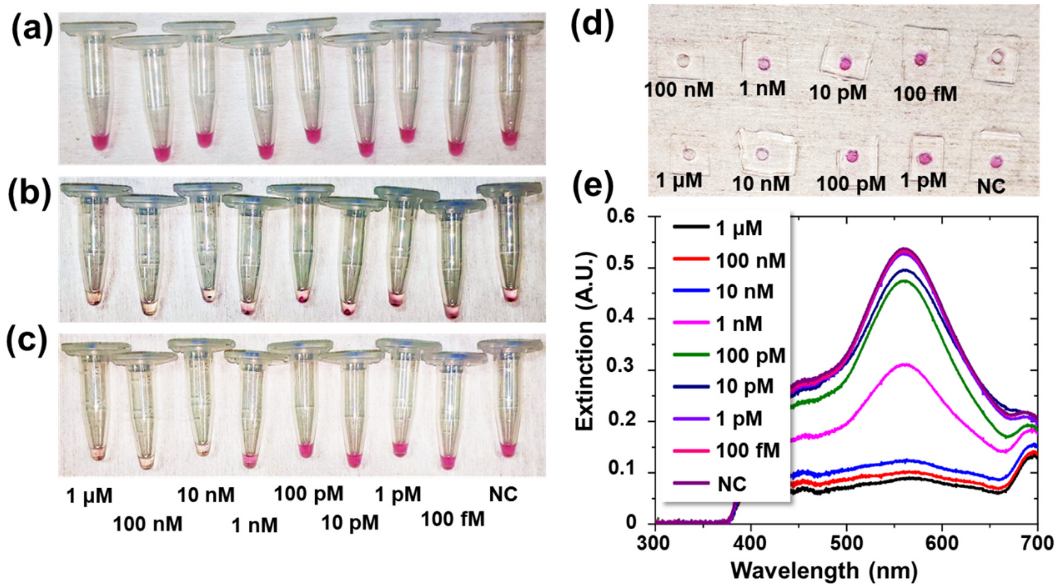


**Figure S18. EBOLA co-binder test in 5% HPS.** (**a-c**) Optical images of microcentrifuge tubes (**a**) after mixing, (**b**) right after centrifugation, and (**c**) after incubation (20 min) and vortex-mixing. (**d**) The upper-level liquid from (**c**) loaded into a PDMS well plate. (**e**) The spectrometric readout from the well plate. The electronic signals shown in Figure 5 were readout from the microcentrifuge tubes after incubation and vortex-mixing.

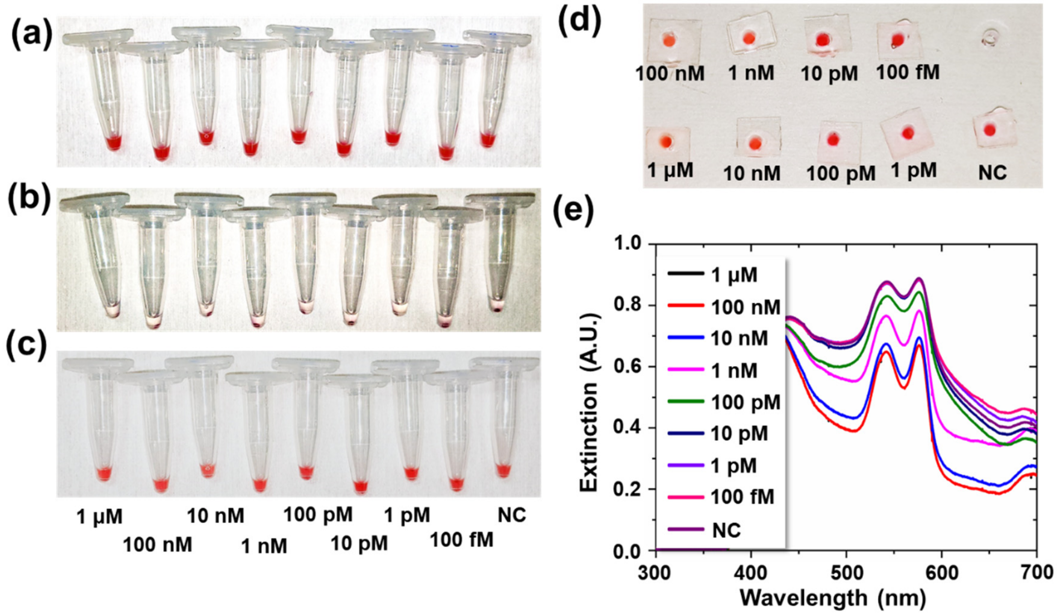


**Figure S19. EBOLA co-binder test in 5% WB.** (**a-c**) Optical images of microcentrifuge tubes (**a**) after mixing, (**b**) right after centrifugation, and (**c**) after incubation (20 min) and vortex-mixing. (**d**) The upper-level liquid from (**c**) loaded into a PDMS well plate. (**e**) The spectrometric readout from the well plate. The electronic signals shown in Figure 5 were readout from the microcentrifuge tubes after incubation and vortex-mixing.

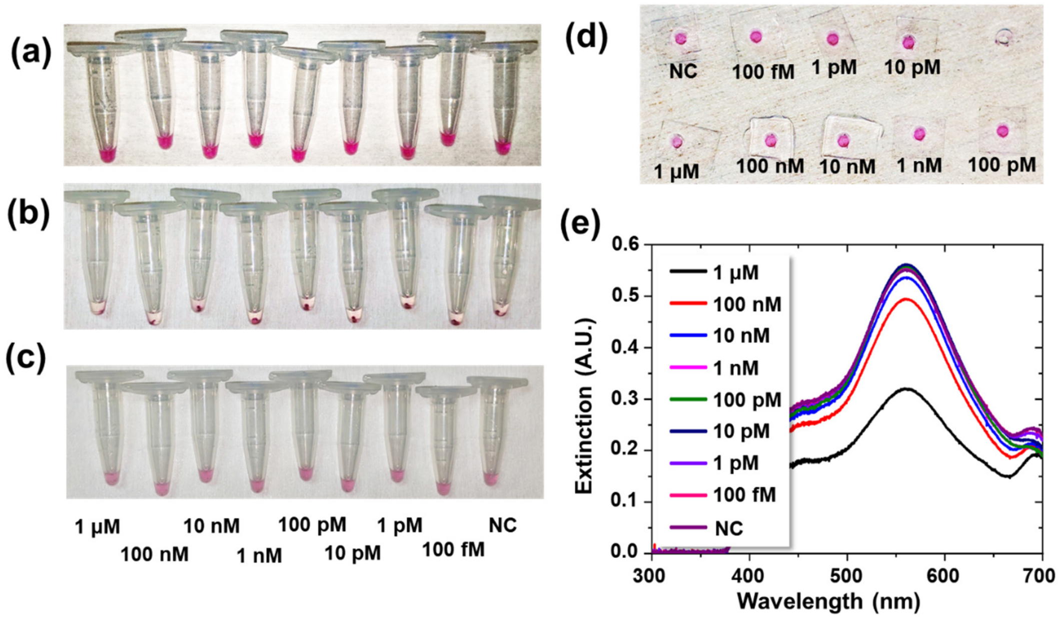


**Figure S20. EBOLA GP1,2 test in 5% FBS.** (**a-c**) Optical images of microcentrifuge tubes (**a**) after mixing, (**b**) right after centrifugation, and (**c**) after incubation (20 min) and vortex-mixing. (**d**) The upper-level liquid from (**c**) loaded into a PDMS well plate. (**e**) The spectrometric readout from the well plate. The electronic signals shown in Figure 5 were readout from the microcentrifuge tubes after incubation and vortex-mixing.

#### 8.3 RBD detection using co-binder RBD8/RBD10

Two different biotinylated nanobodies, RBD10 and RBD8, were surface functionalized to streptavidin coated AuNP similar to the method described earlier for creating two different sets of functionalized AuNP colloidal solutions. The concentration of functionalized AuNP colloids were re-adjusted to get an optimal extinction level. RBD stock solution underwent serial dilution to create an RBD analyte solution with concentrations of 4 uM to 4 pM in selected detection media. The final composition of PBS detection media was composed of 1×PBS, 20% v/v glycerol and 1 wt% BSA while that of FBS, HPS (Human pooled serum), and WB (Whole blood) detection media had a final concentration of 1×PBS, 20% v/v glycerol and 1 wt% BSA and 20% of either FBS, or HPS which resulted in a final concentration of FBS or HPS in the detection assay to be 5%. Solutions of RBD8-functionalized AuNPs, RBD10-functionalized AuNP, and RBD proteins were mixed in a 500 uL Eppendorf tube at a ratio of 3:3:2 and thoroughly vortexed at 800 rpm for 15-20 seconds. After mixing, the detection assay was centrifuged (accuSpin Micro 17, Thermo Fisher) at 3,500 rpm (1,200×g) for 1 minute. After 20 minutes of incubation, the colloidal assay was vortexed (mini vortexer, Thermo Fisher) at 800 rpm for 15 seconds. Then spectrometric and electronic characterizations were performed.

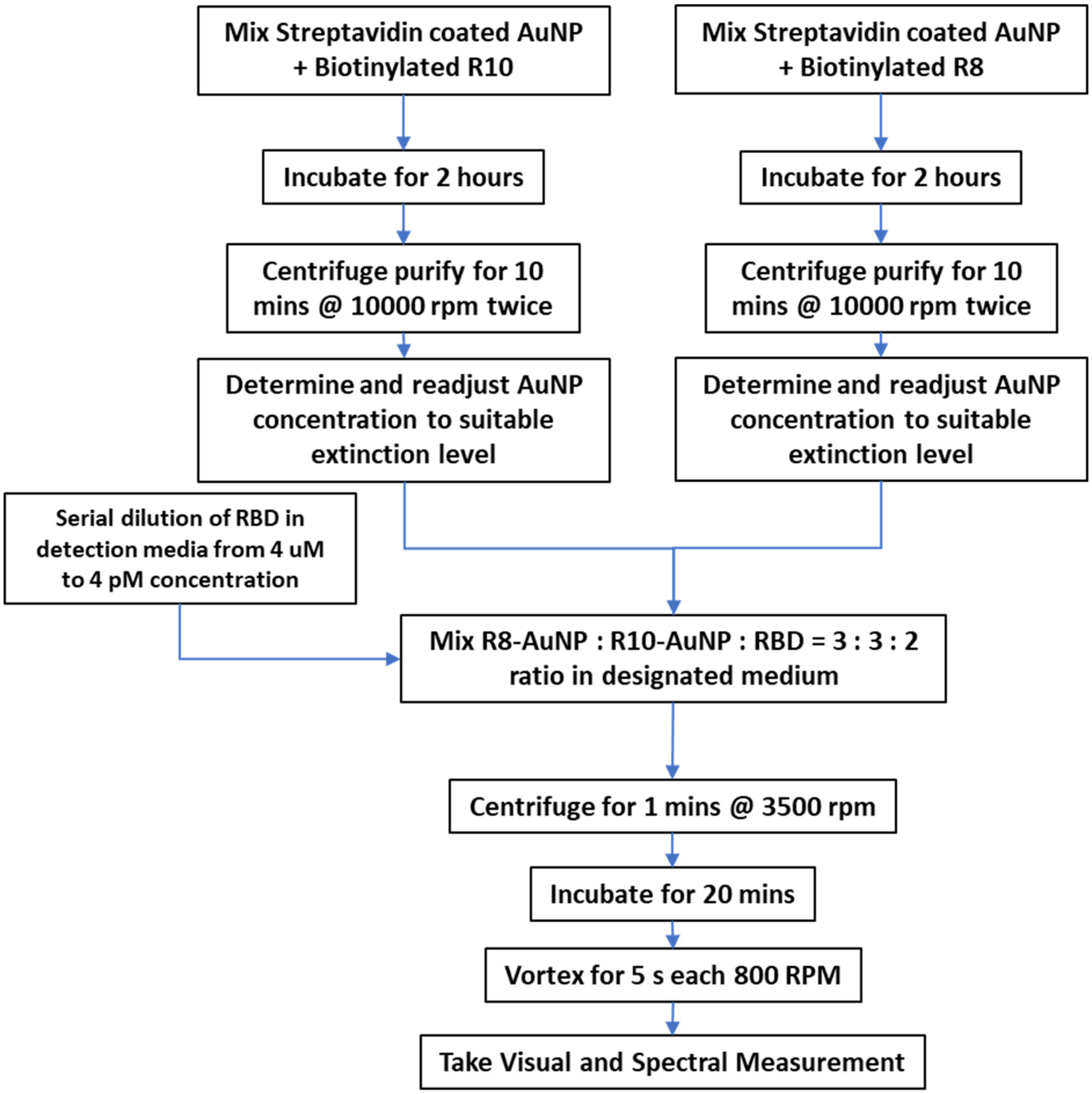


**Figure S21. Workflow of RBD sensing using co-binders.**

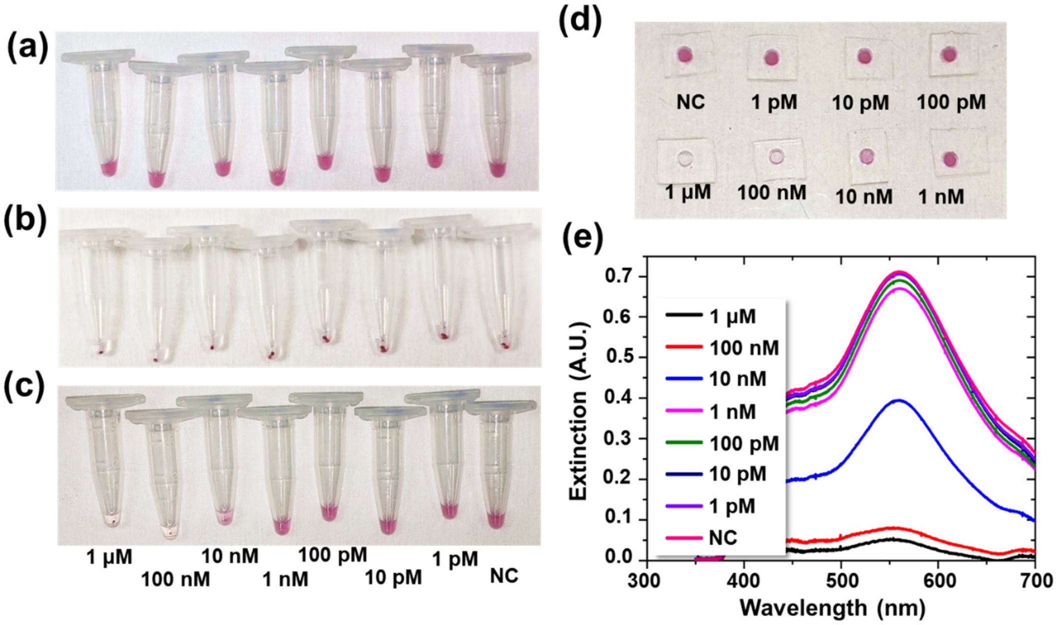


**Figure S22. COVID-19 RBD test in 1X PBS.** (**a-c**) Optical images of microcentrifuge tubes (**a**) after mixing, (**b**) right after centrifugation, and (**c**) after incubation (20 min) and vortex-mixing. (**d**) The upper-level liquid from (**c**) loaded into a PDMS well plate. (**e**) The spectrometric readout from the well plate. The electronic signals shown in Figure 5 were readout from the microcentrifuge tubes after incubation and vortex-mixing.

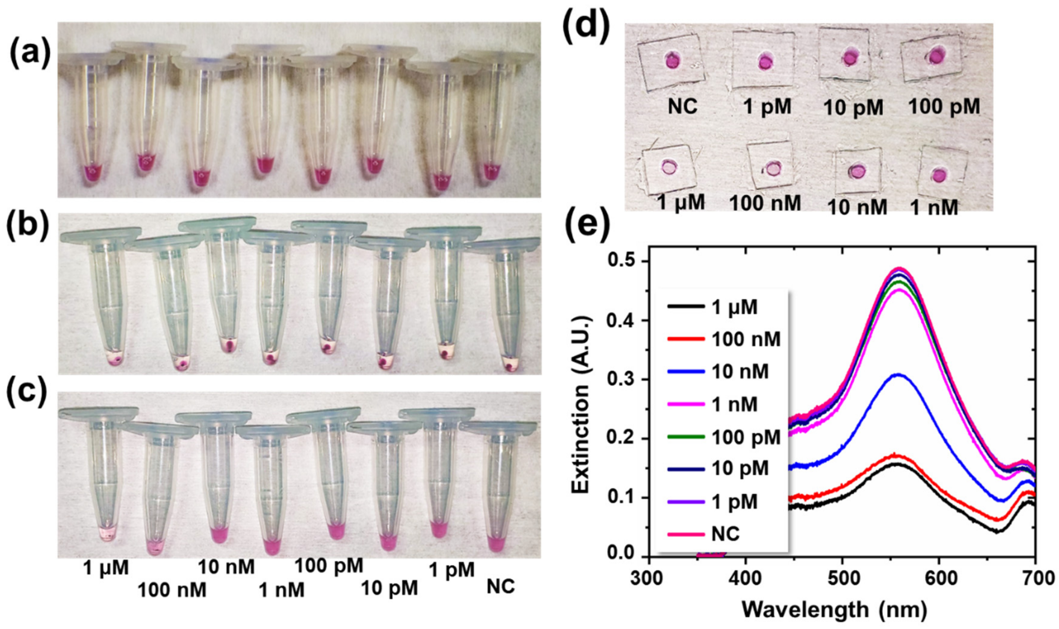


**Figure S23. COVID-19 RBD test in 5%FBS.** (**a-c**) Optical images of microcentrifuge tubes (**a**) after mixing, (**b**) right after centrifugation, and (**c**) after incubation (20 min) and vortex-mixing. (**d**) The upper-level liquid from (**c**) loaded into a PDMS well plate. (**e**) The spectrometric readout from the well plate. The electronic signals shown in Figure 5 were readout from the microcentrifuge tubes after incubation and vortex-mixing.

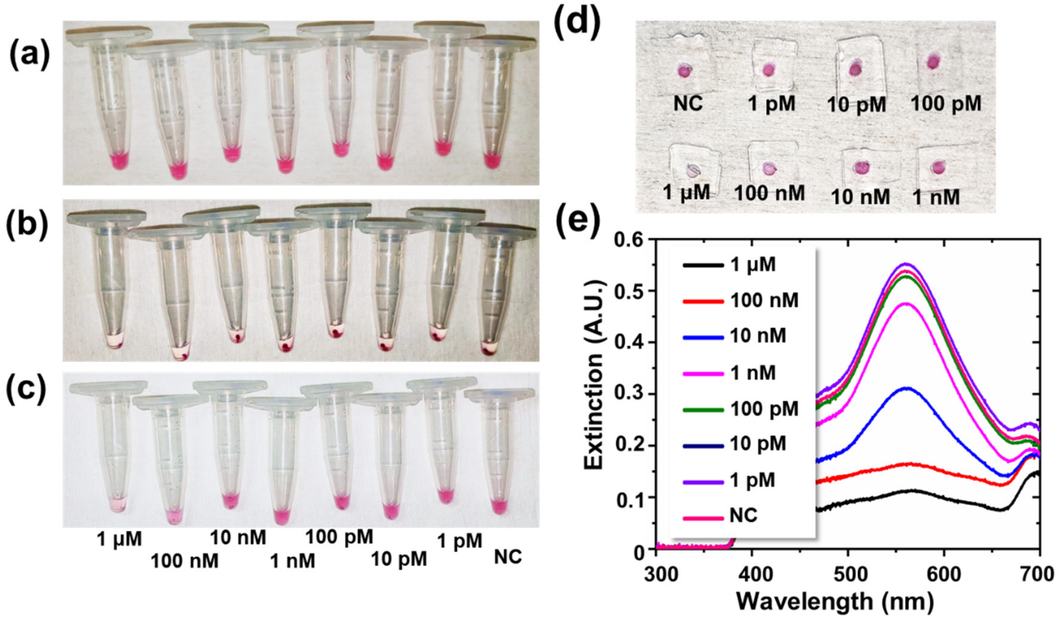


**Figure S24. COVID-19 RBD test in 5%HPS.** (**a-c**) Optical images of microcentrifuge tubes (**a**) after mixing, (**b**) right after centrifugation, and (**c**) after incubation (20 min) and vortex-mixing. (**d**) The upper-level liquid from (**c**) loaded into a PDMS well plate. (**e**) The spectrometric readout from the well plate. The electronic signals shown in Figure 5 were readout from the microcentrifuge tubes after incubation and vortex-mixing.

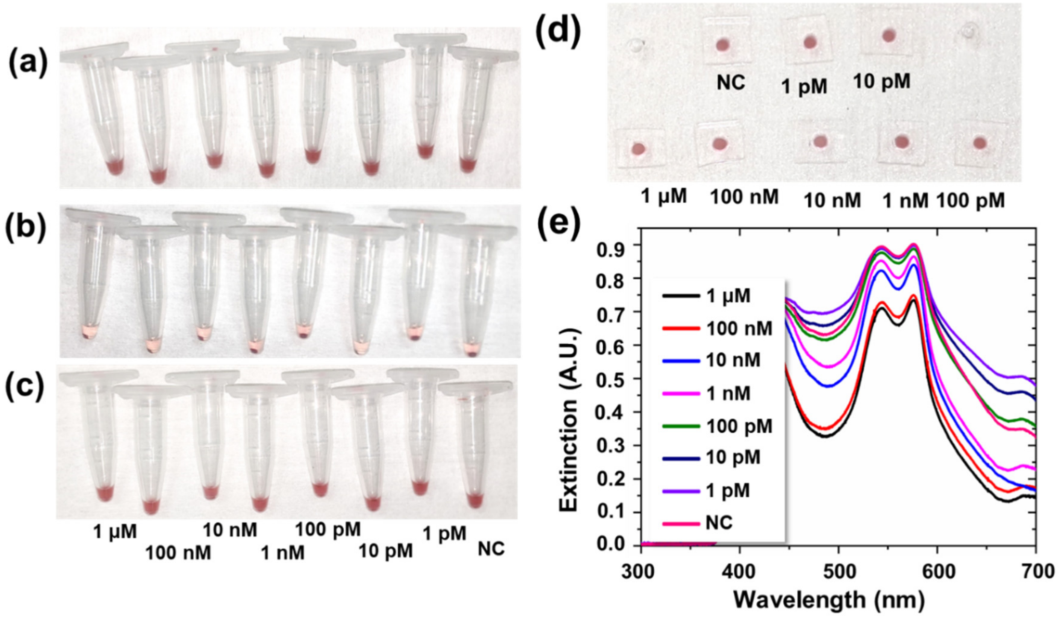


**Figure S25. COVID-19 RBD test in 5%WB.** (**a-c**) Optical images of microcentrifuge tubes (**a**) after mixing, (**b**) right after centrifugation, and (**c**) after incubation (20 min) and vortex-mixing. (**d**) The upper-level liquid from (**c**) loaded into a PDMS well plate. (**e**) The spectrometric readout from the well plate. The electronic signals shown in Figure 5 were readout from the microcentrifuge tubes after incubation and vortex-mixing.

#### 8.4. Detection error and LOD in Phage ELISA

To determine the Phage ELISA LOD analysis similar to the method described in previous section was performed (Table S1). For consistency in LOD anaysis, we analyzed the results with three different methods for sigma estimation. The first method is the traditional way of determining LOD, which only takes into account the standard deviation of blank (or NC) sample measurement (termed $\sigma_{NC}$). The two additional methods, i.e., Pooled Sigma All ($\sigma_{PSA}$) and Pooled Sigma 4 ($\sigma_{PS4}$) employ a pooled variance method which is used for statistical analysis of different populations but with similar variances to give a more robust and consistent result. While $\sigma_{PSA}$ takes into account all of the samples, $\sigma_{PS4}$ takes onto account only the last four samples.

Here for ELISA, the measurement noise is strongly correlated to signal level and sample concentration (Figure S26 and Table S1). As a result, $\sigma_{PSA}$ is quite high due to the consideration of high-concentration samples. Yet the use of $\sigma_{NC}$ makes the result very susceptible to experimental errors in measuring the NC sample, which could seriously affect the LOD by orders of magnitude given a small signal difference at low sample concentration. Comparatively, $\sigma_{PS4}$ provides a more balanced estimation for reagent characterization, because the last 4 sample data (including the NC sample) have similar signal and noise levels and the average in fact provides a more consistent estimation of experimental errors.

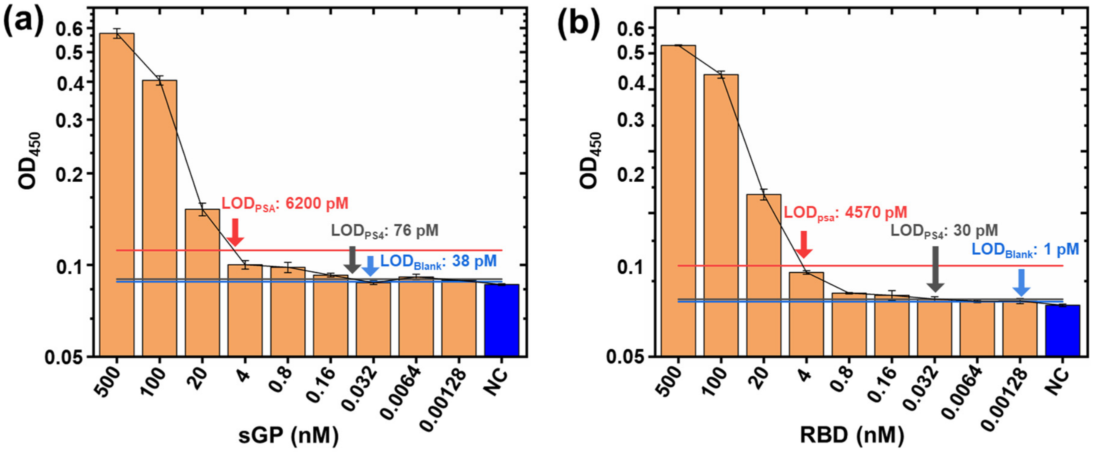


**Figure S26. LOD Measurement in Ebola Cobinder rapid test in PBS.** The extracted $OD_{450}$ values for the Phage ELISA test are plotted with the fitted detection curve (Black). The horizontal lines represent NC-3σ values for Blank sigma (blue), Pooled Sigma 4 (gray) and Pooled Sigma All (red). (a) For sGP detection with mono-binder (sGP49) with LOD of 38 pM, 76 pM and 6200 pM, for Blank, PS4 and PSA respectively, (b) RBD detection with co-binder (R8 and R10) with LOD of 1 pM, 30 pM and 4570 pM, for Blank, PS4 and PSA respectively.

**Table S1. LOD analysis of Ebola sGP and COVID RBD detection with Phage-ELISA**

| Antigen | Binder | Media | Readout method | $\sigma$ (A.U.) | | | LOD (pM) | | |
|---|---|---|---|---|---|---|---|---|---|
| | | | | $\sigma_{PS4}$ | $\sigma_{PSA}$ | $\sigma_{NC}$ | $\sigma_{PS4}$ | $\sigma_{PSA}$ | $\sigma_{NC}$ |
| **Ebola sGP** | sGP49 | PBS | Colorimetric | 0.00121 | 0.00851 | 0.000622 | 76 | 6200 | 38 |
| **COVID-19 RBD** | R8-R10 | PBS | Colorimetric | 0.00415 | 0.0331 | 0.002538 | 30 | 4570 | 1 |

#### 8.5. Detection error and LOD in rapid test

For consistent data acquisition, it was imperative that the spectrometric data acquisition be done as soon as possible in order to minimize error introduced by gravitational precipitation of AuNP. Here it was noticed that the traditional method of determining LOD, i.e. $\sigma_{NC}$, negatively affected the consistency in LOD determination (Table S2). This can be attributed to the nature of data collection in that optical focusing varies from one measurement to the next. In comparison, $\sigma_{PSA}$ and $\sigma_{PS4}$ provide better consistency and robustness in determining the standard deviation required for LOD measurement. This is particularly true when the error measured from the NC sample is too high or too low compared to the average errors measured across the whole sample set. Here, we choose the $\sigma_{PSA}$ method for further analysis of the data because of its better data consistency. Table S1 shows the summary of the analysis while Figure S25 shows the LOD determination using the different calculated SD values.

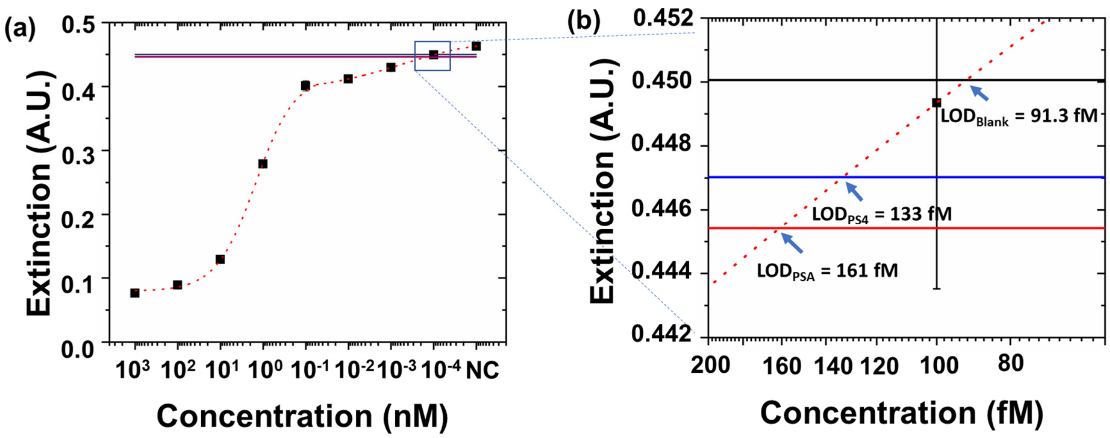


**Figure S27. LOD Measurement in Ebola Cobinder rapid test in PBS. (a)**The extracted extinction peaks at 559 nm are plotted with the fitted detection curve (dotted red line). The horizontal lines represent NC-3σ values for Blank sigma (black), Pooled Sigma 4 (blue) and Pooled Sigma All (red). (b) Zoomed in figure of the area of interest showing the intersections giving the LODs obtained through three different methods with LOD of 91.3 fM, 133 fM and 161 fM, for Blank, PS4 and PSA respectively.

**Table S2. LOD analysis of Ebola sGP and COVID RBD detection by Nano2RED**

| Antigen | Media | Readout method | $\sigma$ (A.U.) | | | LOD (pM) | | |
|---|---|---|---|---|---|---|---|---|
| | | | $\sigma_{PS4}$ | $\sigma_{PSA}$ | $\sigma_{NC}$ | $\sigma_{PS4}$ | $\sigma_{PSA}$ | $\sigma_{NC}$ |
| Ebola sGP | PBS | Spectrometric | 0.005211 | 0.005747 | 0.0042 | 0.133 | 0.161 | 0.0913 |
| | FBS | | 0.007094 | 0.005944 | 0.00914 | 1.65 | 1.05 | 3.75 |
| | HPS | | 0.004836 | 0.00548 | 0.00558 | 1.09 | 1.26 | 1.29 |
| | HPS | Electronic | 0.004459 | 0.004518 | 0.00483 | 0.134 | 0.135 | 0.142 |
| | WB | Spectrometric | 0.005841 | 0.005276 | 0.00701 | 18.24 | 16.9 | 21.44 |
| COVID-19 RBD | PBS | Spectrometric | 0.005325 | 0.00527 | 0.00612 | 1.51 | 1.43 | 3.4 |
| | FBS | | 0.005643 | 0.005621 | 0.00522 | 5.28 | 5.23 | 4.39 |
| | HPS | | 0.00474 | 0.0046 | 0.00346 | 32.93 | 22.25 | 7.29 |
| | WB | | 0.00608 | 0.0087 | 0.00671 | 121.91 | 253.3 | 153.65 |
| | HPS | Electronic | 0.00419 | 0.004288 | 0.00311 | 1.34 | 1.35 | 1.15 |

### 9. Reagent cost analysis

Here, using nanobody-conjugated 80 nm AuNP as an example, we estimated the cost of our AuNP assay, from gold nanoparticle synthesis, nanobody selection, AuNP surface function to final functional assay colloids production.

First, gold nanoparticles are synthesized following standard protocols[11]. Consumption of $HAuCl_4$ ($100/g), Tween 80 ($50/g), Sodium borohydride ($0.4/g), and other additives ($1/g) leads to $110/g estimated cost of bare 80 nm AuNPs. The bare AuNP then undergoes streptavidin surface functioning, involving the consumption of NHS-EDC chemistry ($10/g), streptavidin (estimated $10,000/g), mercapto-succinic acid ($1/g), and working buffers ($5/L). The cost of streptavidin surface functioned 80 nm AuNP is estimated to be $275/g.

Next, the synthetic nanobody is selected following the protocol in the Method section (Phage display selection). The whole process includes the cost from Sanger sequencing ($100), Minipre ($90), Dynabeadse M280 ($200), anti-M13 antibody and second anibody ($20), immune 96-well plates ($10), TMB-ELISA substrate ($15), and SA biosensor ($100). The total cost of nanobody selection is estimated to be $515. Further, the selected nanobody is purified involving yeast extraction ($10), tryptone ($20), isopropylthio-β-galactoside (IPTG) ($1), and imidazole ($1) per 10 mg nanobody. The purified nanobody cost is estimated to be $3200/g. Then, nanobody is biotinylated using BirA ligates ($5), biotin ($0.1) and ATP ($0.1) chemicals per 10 mg nanobody. The final cost for biotinylated nanobody is $3720/g.

Finally, the AuNP colloid assay is made by mixing streptavidin surface functioned AuNP with the nanobodies. To saturate the streptavidin proteins coated on AuNP with nanobodies, each 80 nm AuNP is mixed with 1440 nanobodies. Therefore, each 1g of 80 nm AuNP consumes 7.9 mg nanobody by weight ($29). Further, centrifugation purification is used to extract sGP49 surface functioned AuNP with a yield rate of ~50% following the protocols described in the Method section (Preparation of sGP49 surface functioned AuNP colloid). The cost of sGP 49 surface functioned AuNP is estimated to be \$1894.7 per nanomole AuNP. Assuming each test needs 20 μL 0.05 nM 80 nm AuNPs, the final cost for a 20 μL 0.05 nM assay is ~\$0.002 per test.

In the above estimation, we consider the most important raw materials costs. In reality, the production cost may include other items related to space, electricity and water, as well as fees for using basic instrumentation, e.g., freezers, fridges, centrifuges. Although these are difficult to estimate, we think that the total material cost would still be less than \$0.01 per test for large-scale manufacturing. We also recognize that personnel salary in the production of the materials may contribute to a significant part of the total cost. Such cost would depend, for example, on the scale of production and salary, and thus are difficult to estimate. However, given the use of standard processes in the production of these types of materials, we believe large-scale and automated production of the assays to be feasible.

### 10. Comparison of Nano2RED with representative diagnostic technologies

Table S3. Representative technologies for Ebola diagnostics

| Assay format | Target markers | Diagnostic sensitivity | Readout method | Readout instrument | Assay time | Test cost | Operation complexity |
|---|---|---|---|---|---|---|---|
| rt-PCR[12-16] | Viral RNA | Very high (~$10^2$-$10^4$ copy/mL) | Fluorescence | Large, expensive | Long (~24 hours) | High | High |
| LAMP[17-19] | Viral RNA | Very high (~$10^2$-$10^3$ copy/mL) | Fluorescence; Colorimetric | Large, expensive | Short (<1 hour) | High | High |
| CRISPR[20, 21] | Viral RNA | High (~$10^4$ copy/mL) | Fluorescence; Colorimetric | Portable, low cost | Medium (1-2 hours) | High | High |
| Ag-ELISA[22-24] | Antigen | High (~$10^2$ ng/ml) | Fluorescence | Large, expensive | Long (a few hours) | Medium | Medium |
| Ag-LFA[25] | Antigen | Medium (qualitative) | Colorimetric | Portable, low cost | Short (<1 hour) | Low | Low |
| Ab-ELISA[16, 26] | Antibody | High (~$10^2$ ng/ml) | Fluorescence | Large, expensive | Long (a few hours) | Medium | Medium |
| Ab-LFA[27] | Antibody | Medium (qualitative) | Colorimetric | Portable, low cost | Short (<1 hour) | Low | Low |
| Nano2RED (this work) | Antigen protein | High (~100 fM) | Colorimetric; Electronic | Portable, low cost | Very short (5-20 min) | Low | Low |

Table S4. Representative Ebola sGP protein diagnostic and readout technologies

<table>
<tr><th>Ref</th><th>Research work</th><th>Assay format</th><th>Readout method</th><th>Readout instrument</th><th>Diagnostic sensitivity</th><th>Estimated instrument cost</th><th>Readout system size</th><th>Assay time</th></tr>
<tr><td>28</td><td>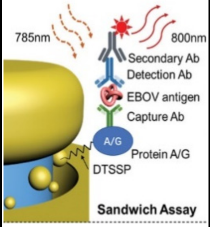
</td><td>Plasmonic sandwich assay</td><td>FI</td><td>Lab microscope with EMCCD camera</td><td>220 fg/mL (2 fM) in human plasma</td><td>>~$40,000 (~ $25,000 for camera)</td><td>~3m$^3$ =3,000,000 cm$^3$</td><td>3-4 hours</td></tr>
<tr><td rowspan="3">29</td><td rowspan="3">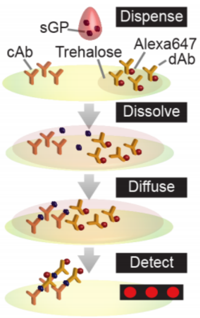
</td><td>D4, co-binder</td><td>FI</td><td>Lab microscope</td><td>20 pg/mL (~ 500 fM) in FBS, 30 pg/mL (~ 750 fM) in HS, 10 pg/mL (~ 250 fM) in WHB</td><td>>~$40,000 (~ $25,000 for camera)</td><td>~3m$^3$ =3,000,000 cm$^3$</td><td>60 min</td></tr>
<tr><td>D4, co-binder</td><td>FI</td><td>Portable fluorescent system</td><td>100 pg/mL (~2.5 pM) in WHB</td><td>$1,000</td><td>21×16 ×9 = 3,000 cm$^3$</td><td>30 min</td></tr>
<tr><td>LFA</td><td>Colori-metric</td><td>Eyes</td><td>6000 pg/mL (150 pM) in FBS/HS/WHB</td><td>~$200 (phone) *</td><td>~30×6 ×0.5 =90 cm$^3$ **</td><td>~60 min</td></tr>
</table>

FI: fluorescent imaging. D4: Dispense, Dissolve, Diffuse, Detect. LFA: Lateral flow assay. HS: Human serum. FBS: Fetal bovine serum. WHB: Whole human blood.

* We assume phones for digitizing the data. For naked eye readout, the cost is close to zero.

** We assume typical dimensions for a lateral flow assay.
